# Patches: A Representation Learning Framework for Decoding Shared and Condition-Specific Transcriptional Programs in Wound Healing

**DOI:** 10.1101/2024.12.23.630186

**Authors:** Ozgur Beker, Simon Van Deursen, Michel Tarnow, Dreyton Amador, Jonathan Chin Cheong, Jose Francisco Pomarino Nima, Mark D. Robinson, Yvon Woappi, Bianca Dumitrascu

## Abstract

Single-cell genomics enables the study of cell states and cell state transitions across biological conditions like aging, drug treatment, or injury. However, existing computational methods often struggle to simultaneously disentangle shared and condition-specific transcriptional patterns, particularly in experimental designs with missing data, unmatched cell populations, or complex attribute combinations. To address these challenges, *Patches* identifies universal transcriptomic features alongside condition-dependent variations in scRNA-seq data. Using conditional subspace learning, Patches enables robust integration, cross-condition prediction, and biologically interpretable representations of gene expression. Unlike prior methods, Patches excels in experimental designs with multiple attributes, such as age, treatment, and temporal dynamics, distinguishing general cellular mechanisms from condition-dependent changes. We applied *Patches* to both simulated data and real transcriptomic datasets from skin injury models, focusing on the effects of aging and drug treatment. *Patches* revealed shared wound healing patterns and condition-specific changes in cell behavior and extracellular matrix remodeling. These insights deepen our understanding of tissue repair and can identify potential biomarkers for therapeutic interventions, particularly in contexts where the experimental design is complicated by missing or difficult-to-collect data.

## Introduction

All living organisms have the remarkable ability to repair themselves in response to injury. Healing efficiently and successfully is essential to preserving the integrity of tissues and organs. As a central biological process, wound healing involves the complex, coordinated actions of numerous intracellular and intercellular components. These include the activation and regulation of different cell types, such as fibroblasts, keratinocytes, and immune cells, as well as the interplay of signaling molecules[1, 2, 3]. Single-cell RNA sequencing (scRNA-seq) provides a detailed view of the transcriptional dynamics during tissue repair, particularly in murine skin wounds [4, 5, 6, 7, 8]. These recent studies have highlighted the heterogeneity of cell populations involved in the process, identifying distinct wound healing processes linked to physiological changes like aging [4], and demonstrating unique contributions of epithelial [5],[9], mesenchymal [6, 7], and immune cells [8] at the injury site. However, despite advances in data collection, much of the current work focuses on selecting individual genes based on prior characterization, limiting our ability to uncover novel regulatory programs. As a result, a systematic understanding of the molecular changes of diverse skin cell lineages during the healing process remains incomplete [10, 9]. This gap highlights the need for new computational tools capable of *de novo* discovery of cell-specific gene programs from complex scRNA-seq wound datasets, enabling an unbiased exploration of the molecular mechanisms underlying repair mechanisms.

Traditionally, scRNA-seq analysis methods have focused on static snapshots of cellular states, which do not adequately capture the dynamic nature of injury response and result in missing or incomplete data. Indeed, the biological complexity and physiological variability of skin tissue repair poses unique challenges[9, 11]. High dropout rates in scRNA-seq, coupled with destructive experimental designs, result in incomplete data and biases that obscure transcriptional dynamics across conditions and timepoints. Furthermore, conditions such as aging, drug treatment, or disease can alter these trajectories in ways that are not fully understood. Many gene programs are shared across conditions, making it statistically challenging to disentangle common mechanisms from those that are condition-specific. Addressing these challenges requires new models capable of integrating longitudinal data, accurately capturing shared patterns, and isolating differences unique to each condition.

A wide range of statistical machine learning methods have been developed to annotate [12, 13, 14], integrate [12, 13, 15, 16], and interpret [17, 18, 19] scRNA-seq data. These approaches differ in their statistical focus whether they: a) are generative or non-generative, b) can capture shared versus condition-specific gene expression patterns in multi-condition datasets, and c) are interpretable. In particular, non-generative approaches have provided pipelines for unsupervised clustering [12, 13, 14], uncovering gene expression programs through matrix factorization techniques [17, 18] and understanding cell-cell interaction [20, 21, 22]. Generative models use probabilistic frameworks to represent scRNA-seq data as nonlinear transformations of samples from a lower-dimensional latent space. Unlike non-generative models, the latent representations learnt by generative models have proven useful in modeling complex biological and technical variation [16, 23, 24, 25, 26]. Additionally, generative models have also been used to integrate data from multiple modalities [27, 28] or conditions [25, 29, 30, 31]. In doing so, these tools focus on either modeling shared signals across all conditions [16, 24, 26] or on modeling conditional information – discrete (e.g., cell type labels [25, 29, 30, 32]) or continuous (e.g., for visualization [23, 31]). However, these methods do not quantify the trade-off between the information shared across all conditions and that is specific to each individual condition. Only a few studies have addressed this challenge [33, 34, 35, 36] particularly when integrating multi-modal data, such as morphology and gene expression [33, 34] or gene expression and chromatin accessibility [35, 36]. These approaches generally require matched cells across modalities, which limits their applicability to destructive and complex experimental designs where the focus is on capturing temporal variation. Still, generative models can capture nonlinear relationships, making them valuable for studying the complex dynamics of wound healing. However, their expressivity often comes at the cost of interpretability compared to non-generative methods. Recent efforts have addressed this trade-off by either incorporating linear layers in the generative process to encourage interpretability [19] or applying linear models post hoc to connect learned representations with generated gene expression profiles [37]. Specifically, the former approach can be used in tandem with conditional generative model formulations to enable interpretability in complex contexts. Taken together, these considerations highlight the need for interpretable generative models specifically designed to decouple shared and condition-specific effects, enabling the study of intricate processes such as wound healing.

To this end, we introduce *Patches*, a deep representation learning framework designed to model gene expression data from complex, multi-condition experimental designs. Similar to existing generative models such as scVI [16, 24, 26], *Patches* maps a cell’s gene expression profile to a low-dimensional latent representation via learnable, nonlinear transformations. However, unlike these approaches, *Patches* leverages conditional subspace learning [38, 39] to disentangle shared and condition-specific transcriptomic patterns. This enables the identification of both common and distinct gene expression features across conditions such as age, treatment type, or temporal profile. This flexible latent space, also allows the generation of transcriptomic data for previously unseen condition combinations by simply modifying the condition labels during generation. By applying *Patches* to both simulated and real datasets, we show it can uncover new insights into the molecular dynamics of wound repair. Specifically, we apply *Patches* to single-cell data from skin injury experiments, demonstrating its ability to capture both universal and condition-dependent dynamics in the contexts of aging and drug treatment. In aging, *Patches* identifies subtle changes in immune cell behavior and extracellular matrix (ECM) remodeling that distinguish young and aged responses to injury. Similarly, in a drug treatment context, *Patches* captures transcriptional changes associated with wound healing dynamics under verteporfin treatment, revealing potential pathways for therapeutic intervention. These results highlight the potential of *Patches* to provide insights into complex cellular processes and inform experimental designs in challenging multi-condition settings.

## Results

### Patches

#### Overview

*Patches* takes as input a collection of observations or gene expression profiles obtained from multiple conditions (Fig. 1A) referred to as ‘groups’ – for example cell type, age, treatment or injury stage. Each group is characterized by a unique set of categories or attributes – for example, ‘young’ or ‘old’ represent the attributes of the group ‘age’.

**Figure 1:**
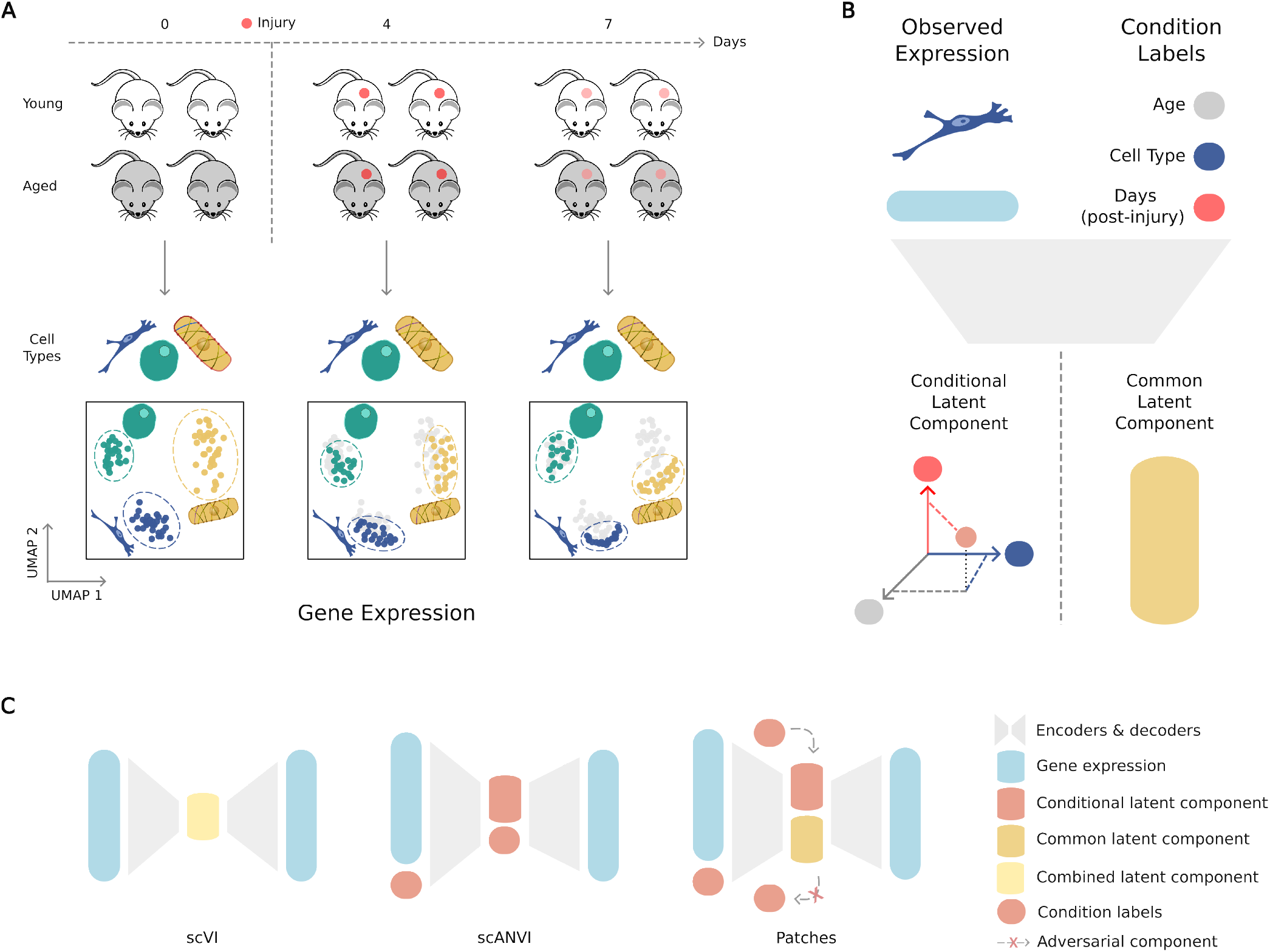
Overview of *Patches* for multi-condition experimental designs: **A**, Graphical illustration of datasets compatible with *Patches*, with observations from different time, age or cell type attributes. **B**, Illustration of how *Patches* models multi-condition data, disentangling condition-specific and condition-agnostic (common) representations. The condition-specific representation captures the expression components that vary across attribute labels, while the common representation reflects shared features across all conditions. **C**, Comparison of *Patches* with existing models: in contrast to scVI, which learns representations in a condition agnostic fashion, and to scANVI, which learns conditional-representations one condition at the time, *Patches* employs a mix of both approaches, obtaining both conditional and common representations, which are then disentangled using information-theoretical or adversarial priors.

Extending existing variational autoencoder (VAE) architectures [16] by incorporating concepts from multi-modal learning [40] and information theory [38, 39], *Patches* models a gene expression profile **x**_*n*_ as a sample from a distribution conditional on a latent representation *ρ*_*n*_ – the cell’s ‘latent identity’. The latent identity *ρ*_*n*_ is decomposed into two components: **z**_*n*_ and **w**_*n*_. This decomposition reflects the interaction between the cell’s intrinsic state and the experimental conditions or attributes under which it was observed. The shared latent variable **z**_*n*_ captures condition-independent features of the cell, remaining invariant across different combinations of experimental attributes. Conversely, **w**_**n**_ encodes group and attribute-specific information that varies according to experimental factors such as age, treatment, or temporal stage (Fig. 1B). This positions *Patches* as a middle ground between unsupervised methods like scVI [16], which focus on learning shared latent spaces, and supervised approaches like scANVI [25], which emphasize condition-specific representations. In contrast to these methods, *Patches* jointly learns both shared (common) and condition-specific embeddings (Fig. 1C). *Patches* also enforces information-theoretic constraints [38, 39] to ensure that the shared embeddings remain disentangled from condition-specific attributes. The constraints are enforced using adversarial classifiers that encourage the shared latent variables **z**_*n*_ to carry minimal information about conditions. In detail, the group-specific latent representations **w**_*n*_ are obtained by concatenating latent representations corresponding to individual group combinations 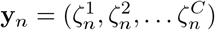 (Fig. 2A). Here, *C* denotes the number of groups represented in the experimental design, for example *C* = 3 when the groups considered are the cell type identity, the age of the sample – whether it is from an old or young mouse – and the wound healing stage (*Methods*). Following previous work, the gene expression *x*_*ng*_ of a gene *g* and cell *n* is modeled using a zero-inflated negative binomial (ZINB) *p*(*x*_*ng*_ | *ρ*_*n*_, *ℓ*_*n*_) [16, 41, 42, 43]. Here, *ℓ*_*n*_ is a cell-specific library size factor and the parameter *ρ*_*n*_ is drawn from a multivariate normal distribution with mean and covariance parameterized by a non-linear function of the latent identities **z**_*n*_ and **w**_*n*_ (Fig. 2A). Finally, while *Patches* inherits the inherent opacity of VAE-based models in its learned representations, it addresses this limitation through an interpretable decoder design. By utilizing a single linear layer [19], *Patches* learns interpretable coefficients that connect original data features, such as gene expression values, to condition-specific attributes. These interpretable coefficients provide a direct explanation of how specific genes contribute to observed differences across experimental conditions (*Methods*). This dual capability of disentangling shared and condition-specific signals while offering interpretable insights makes *Patches* a robust and versatile tool for analyzing complex scRNA-seq datasets.

**Figure 2:**
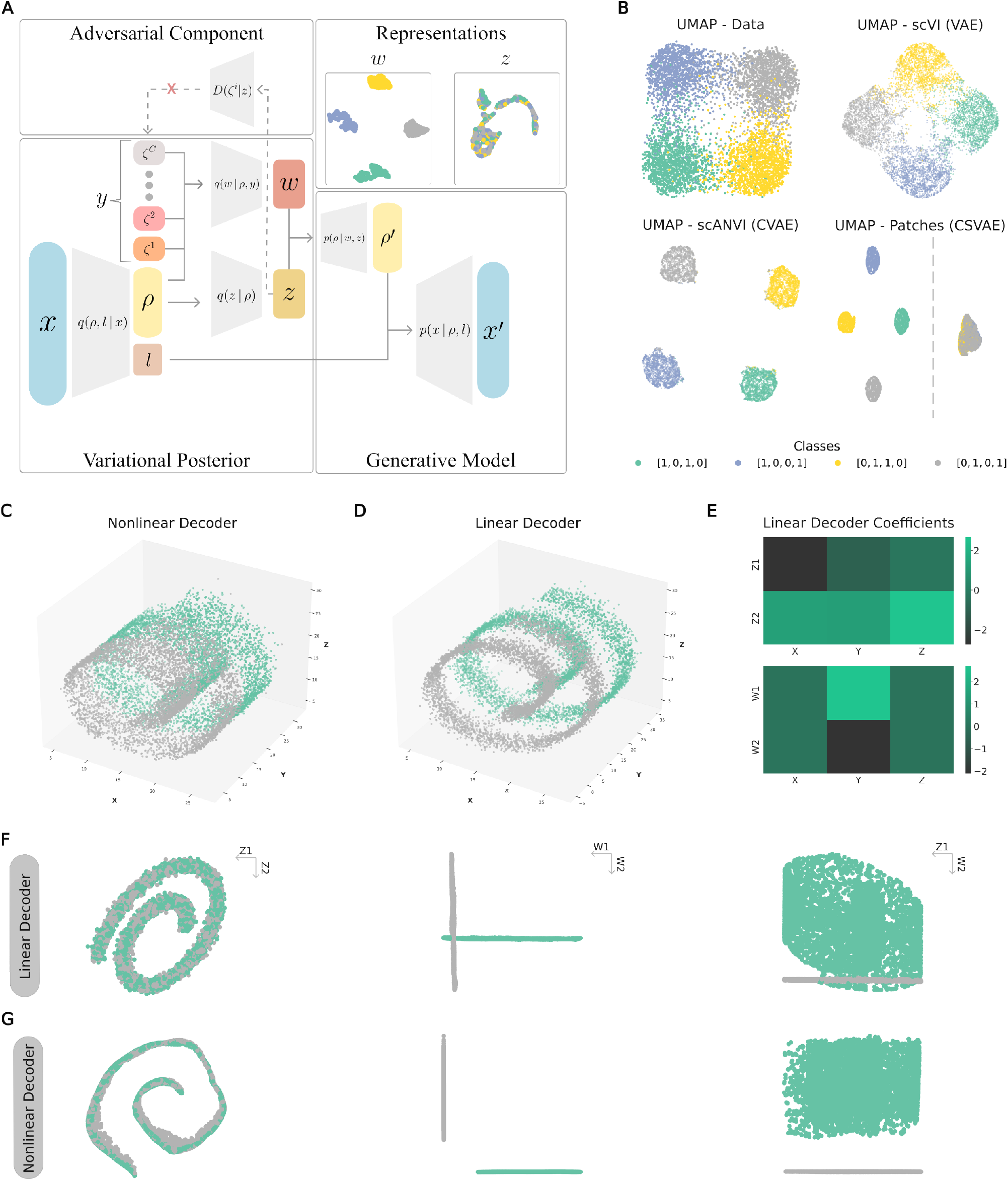
A proof of concept for *Patches* using synthetic examples. Given observations with conditional labels as input, *Patches* obtains condition-specific and common representations. **A**, Graphical model for *Patches*: the adversarial discriminators *D*(*ζ*^*i*^ | **z**) and all the approximate distributions are implemented via neural networks. **B**, UMAP embeddings of the original observations and their corresponding latent representations for the synthetic attribute separation model, demonstrating the ability of *Patches* to separate attributes. **C, D**, Reconstruction of the Swiss roll by *Patches* with nonlinear (**C**) and linear (**D**) decoders, showing differences in reconstruction quality. **E**, The linear decoder in *Patches* learns coefficients that indicate the contribution of each variable in the original data space (columns) to the latent space (rows). **F**,**G** Latent representations of the Swiss roll data for linear (**F**) and nonlinear **G** decoders, highlighting the separation of common and condition-specific components. Legends in the top right of the panels indicate axes.

### Illustrative examples

As proof of concept, we applied *Patches* to two synthetic datasets, comparing its performance with other generative modeling approaches and highlighting its capacity to separate universal patterns from those tied to specific experimental conditions (*Methods*).

#### Synthetic attribute separation model

We first simulated a toy dataset of *n* = 8, 000 data points, each represented by a *G*-dimensional feature vector *x*_*n*_ ∈ ℝ^*G*^, with *G* = 30 (Fig. 2B). The dataset was designed to mimic an experimental setup with two condition groups, *ζ*^1^ and *ζ*^2^, each defined by two attributes. The labels, 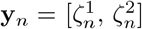, were obtained by concatenating the one-hot encoded attributes of each group, resulting in four possible configurations: [1, 0, 1, 0], [1, 0, 0, 1], [0, 1, 1, 0], and [0, 1, 0, 1] (*Methods*). We simulated 2, 000 points from each configuration as follows. We sampled latent representations 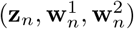 consisting of three concatenated vectors: a shared component **z**_*n*_ ∈ ℝ^10^ and two condition-specific components, 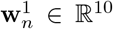 and 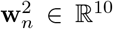. The shared component **z**_*n*_ was sampled from a multivariate normal distribution with mean **0**_10_ and covariance 0.5 𝕀_10_, where **b**_10_ denotes a vector of size 10 with entries all equal to a constant value *b*. The conditional latent component 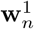 was drawn from a multivariate normal distribution with covariance 0.5 𝕀_10_ and mean **1.15**_10_ (resp. **0**_10_) for *ζ*^1^ = [0, 1] (resp. *ζ*^1^ = [1, 0]). Similarly, 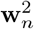 was drawn conditionally from a multivariate normal distribution with covariance 0.5 𝕀_10_ and mean −**1.15**_10_ (resp. **0**_10_) for *ζ*^2^ = [0, 1] (resp. *ζ*^2^ = [1, 0]). Then, the observations **x**_*n*_ were conditionally sampled given latent representations 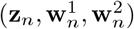 from a multivariate normal distribution with mean 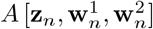 and covariance 0.5 𝕀_30_, with *A* the identity matrix, corresponding to a linear, non-compressive decoder.

We found that *Patches* correctly decomposed each simulated synthetic profile into attribute-specific components and a shared component (Fig. 2B). Specifically, each data point was projected together with points that were originally clustered in the conditional space (left) and onto a shared cluster in the condition-agnostic or common space (right). In this synthetic scenario, scANVI [25] captures only attribute-specific representations, while scVI [16] biases these representations toward a shared component, failing to accurately model either joint or common behavior. In contrast, Patches effectively disentangles shared and condition-specific variation.

#### Supervised Swiss roll

We also applied *Patches* to a popular machine learning toy-data set, the *Swiss roll* [44]. This synthetic dataset is often used to illustrate nonlinear structures in high-dimensional data: it represents a 2D plane curved into a 3D spiral, where local relationships between points are preserved, but global distances can be misleading due to the nonlinearity of the embedding (Fig. 2C). VAEs have been previously used on the Swiss roll to show their capacity in learning nonlinear data, ‘unrolling’ the Swiss roll into flattened latent representations [38]. We generated *n* = 10, 000 points in the shape of a Swiss roll (each point has coordinate features *X, Y, Z*), using an existing implementation [44, 45]. We sliced the Swiss roll structure along the *Y*-axis into two equal parts at *Y* = 10, and labeled the points based on their slice of origin. The resulting supervised Swiss roll dataset contained a single binary condition (*C* = 1, *K* = 2) representing the two slices. We trained *Patches* on this labeled dataset using latent dimensions *d*_**z**_ = 2 and 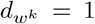 (*Methods*), where each latent dimension *w*^*k*^ corresponds to one of the slices. We analyzed two configurations of *Patches*: one with a nonlinear decoder and another with a linear decoder.

As expected, the nonlinear decoder outperformed the linear decoder in reconstruction performance, successfully capturing the full surface of the Swiss roll (Fig. 2C). In contrast, the linear decoder struggled to reconstruct large portions of the data (Fig. 2D). However, the linear decoder compensated for its limited expressivity with greater interpretability. Unlike the black-box nature of the nonlinear decoder, the linear decoder’s weights provide insights into the relationship between the original data features and their labels (Fig. 2E). Specifically, the decoder coefficients for the latent dimensions *w*^1^ and *w*^2^ clearly distinguish whether a slice is located in the front or back of the Swiss roll based on the Y-coordinate, when viewed from a fixed angle. Further, the model did not wrongly associate any of the other dimensions (*X, Z*) with labels (Fig. 2E). Notably, both the linear and non-linear configurations of *Patches* learned latent representations that reflect the underlying properties of the data-generating mechanism (Fig. 2F,G). Particularly, the shared latent components captured the global structure of the Swiss roll (Fig. 2F,G; left panels), while the condition-specific components accurately represent the binary *Y*-conditions (Fig. 2F,G; middle panels). Finally, the information-theoretic constraint encouraged the latent spaces to remain disjoint, ensuring a clear separation of shared and condition-specific variations (Fig. 2F,G; right panels).

#### Embedding-based gene scoring recovers cell-type and condition specificity in a simulation study

To clarify what is gained by working with embeddings learned by *Patches*, we compare two approaches for gene scoring under the same simulation and evaluation framework. The first scores genes directly using expression-level summary statistics, as in a related and identifiable feature-selection framework [46], where genes are assigned typeness and stateness scores reflecting their association with cell identity and experimental condition. The second derives analogous scores from a learned low-dimensional embedding of cells. In the feature-based framework, typeness and stateness scores are computed from a collection of expression-level statistics (Sup. Tab. 1) and used to define gene-selection strategies (Sup. Tab. 2). Rather than introducing new summary statistics, we ask whether embeddings learned by *Patches* support the same decomposition. To this end, we define corresponding typeness- and stateness-like scores directly in the *Patches* embedding space (Sup. Tab. 3) and use them to construct comparable gene-selection strategies (Sup. Tab. 4). This design isolates the contribution of the learned representation while keeping the downstream scoring and evaluation criteria aligned. We evaluate this comparison using the simulation setup of Wang et al. [46], generated with *Splatter* [47], which includes three cell types and two conditions (Sup. Fig. S1). The task is to recover the ground-truth sets of genes associated with cell type and condition using the same metrics for both approaches. We find that gene scores derived from the *Patches* embedding recover the same type- and condition-associated genes identified by feature-based statistics, particularly for mean absolute deviation–based measures (Sup. Fig. S2). Together, these results indicate that once the embedding is learned, the relevant gene-level effects can be recovered using a linear decoder, without the need for additional complex nonlinear models.

Taken together, these results suggest that both versions of Patches learn informative latent representations that disentangle conditional variation from common signals. However, their distinct strengths make them suitable for different tasks. The nonlinear decoder’s reconstruction performance makes it a good candidate for applications like transfer learning or imputation. These tasks require inferring missing data or generating synthetic data for scenarios where collection is difficult. In contrast, the linear decoder offers greater interpretability. Hence, it is better suited for hypothesis generation, where understanding and acting on the signal is key. For example, it can identify gene programs that are condition-agnostic (e.g., metabolic processes essential for normal function) or condition-specific (e.g., wound-healing pathways activated in response to injury or hypoxia-induced signaling during low oxygen conditions).

### *Patches* learns shared and attribute-specific representations in wound healing studies

Next, we demonstrated the performance of *Patches* on two datasets with multi-group experimental designs. The first dataset investigates the impact of aging on wound healing dynamics [4], while the second explores the effects of drug treatment [48]. These datasets included labels capturing contextual conditions, such as experimental timepoints or treatments, as well as intrinsic identities, such as cell types.

#### Aging

We first applied *Patches* to the wound healing dataset consisting of 26, 966 cells collected across three timepoints (0DPW, 4DPW, 7DPW) from young (7-week-old) and aged (88-week-old) mice [4]. In addition to gene expression profiles, the model was provided conditional information on time post-injury, age, and cell type annotations. *Patches* learned both common (*z*) and condition-specific (*w*) representations for each cell (Fig. 3A). In the conditional latent space, UMAP visualizations revealed distinct clusters corresponding to each combination of attributes (e.g., 4DPW young fibroblasts), consistent with the behavior observed in simulations. The common representations (*z*) effectively integrated data across conditions, aligning cells from different groups, underlying gene expression programs which are common to the wound healing process regardless of age. We further visualized the first two principal components or PCs obtained from the latent gene expression profiles corresponding to shared and condition-specific signals, respectively (Fig. 3A, last column). This confirmed that attribute separation occurred exclusively in the conditional latent space.

**Figure 3:**
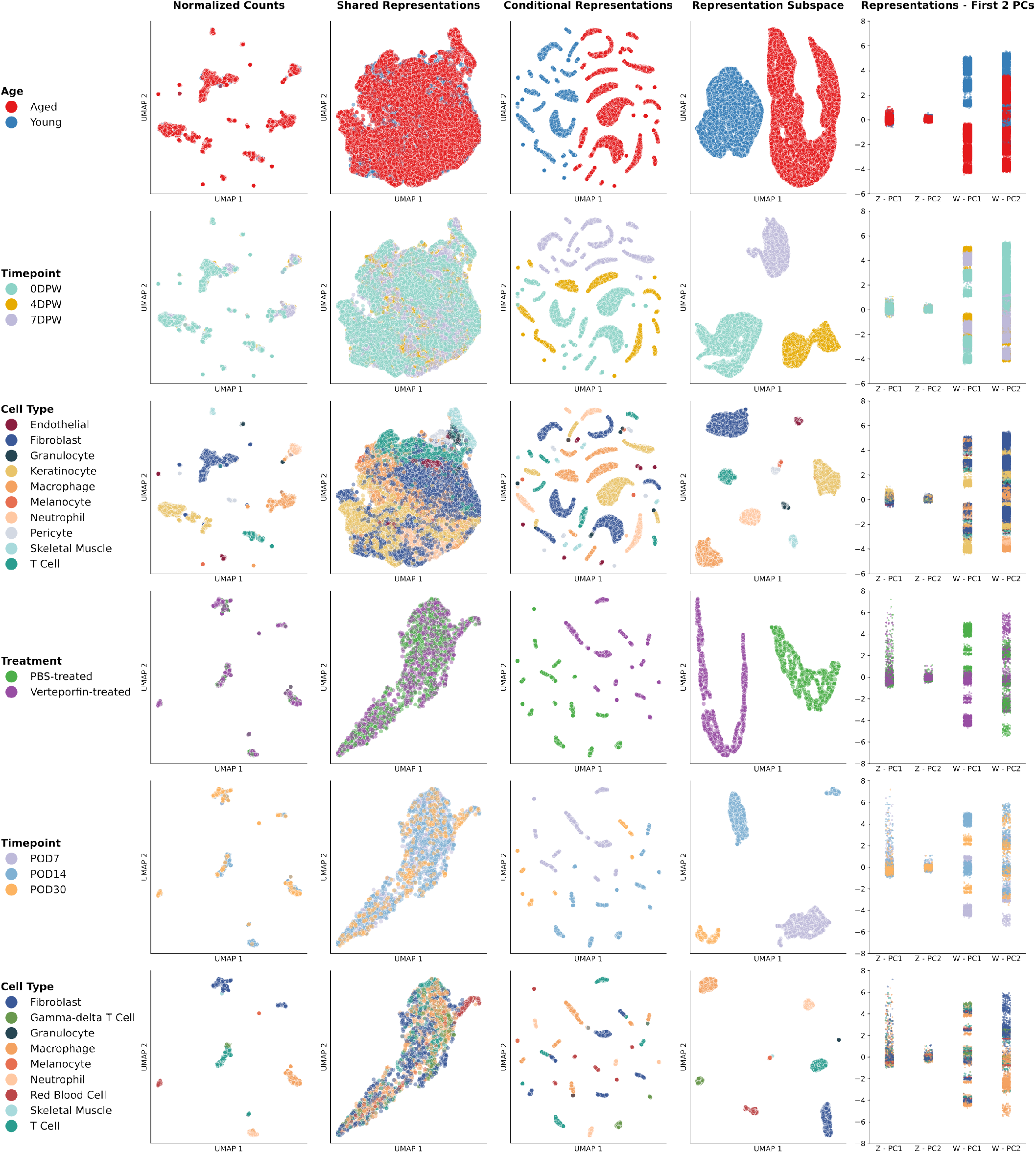
Application of *Patches* to wound healing datasets. **A, B**, Visualization results for gene expression profiles from aged and young mice, collected from 0 (unwounded), 4 and 7 days post wounding conditions (**A**) [4] and for gene expression profiles collected from mice 7, 14 or 30 post operative days (POD), treated with either PBS (control) or Verteporfin **B** [48]. Cells were colored by condition groups across rows, corresponding to cell types (**A, B**) time (**A, B**), age (**A**) and treatment (**B**). The first four columns show UMAP embeddings of: the original gene expression profiles (first), the common latent representations (second), the conditional latent representations (third), and conditional subspace representations (fourth). The fifth column shows the values of the first two principal components, with the first and second inner columns corresponding to common latent representations, and the third and fourth inner columns corresponding to conditional latent representations.

#### Drug treatment

To further evaluate the performance of *Patches*, we analyzed a dataset examining the effect of drug treatment on wound healing [48]. This dataset posed a specific challenge common to many multi-conditional datasets: a significant imbalance in the number of cells per condition (cell type, wound healing stage, drug treatment). After preprocessing, the dataset consisted of 2, 284 cells collected at three timepoints (POD7, POD14, POD30) from either PBS (control) or verteporfin treated mice. As before, for each cell, the model was provided with the time post-injury, treatment, and cell type labels alongside the corresponding gene expression profiles. Despite the low observation count, *Patches* successfully learned common (*z*) and condition-specific (*w*) representations for each observation (Fig. 3B). Unlike in the aging dataset, the low observation count resulted in more sparsely populated clusters in the conditional space, each corresponding to a unique combination of attributes (e.g., POD7 verteporfin treated fibroblasts). Notably, as with similar methods [16, 19], the low dataset size required an increase in the number of training iterations to achieve convergence, as parameter updates depend on dataset size. This observation aligns with previous studies tackling low-sample scenarios (*Methods*).

### Quantifying the attribute-specific representations learned by *Patches*

In both the aging study [4] and the drug treatment study [48], *Patches* learned latent representations that could be qualitatively associated with attributes through UMAP visualizations (Fig. 3). However, visualization alone does not provide a quantitative measure of performance, nor does it explain how these representations reflect the underlying biological attributes. To address this, we further investigated the latent representations learned by *Patches*. Motivated by the results from the simulated Swiss roll example, we evaluated the representations in both linear and nonlinear scenarios. Specifically, we examined *Patches*’ interpretability module and quantified the latent space separability between attribute-specific representations. These were benchmarked against latent spaces learned by previously proposed generative models.

#### *Patches* recovers cell type markers

To test its capacity for extracting interpretable signals, we applied *Patches* with a linear decoder to real scRNA-seq wound healing datasets. [4, 48]. Alongside gene expression data, we incorporated auxiliary annotations, including time and age labels for the aging dataset, time and treatment labels for the drug treatment dataset, and cell type information. After model training, we derived attribute-specific scores for each gene using the learned coefficients from the linear decoder (*Methods*).

We analyzed cell type-specific scores, focusing on the top 20 genes for the three predominant cell types in each dataset. In the aging dataset, these included fibroblasts, keratinocytes, and macrophages (Fig. 4A), while in the drug treatment dataset, they comprised macrophages, fibroblasts, and T cells (Fig. 4B). *Patches* accurately linked key marker genes to their respective cell types. For instance, fibroblast markers such as *Dcn, Col1a1, Col1a2*, and *Col3a1* were identified (Fig. 4A,B; top row), along with keratinocyte markers, including *Krt14* and related *Krt* genes (Fig. 4A, middle row). For macrophages, *Lyz2* and *Cd14* were recovered (Fig. 4A, bottom row; Fig. 4B, middle row), while T cell markers included *Nkg7* and *Ccl5* (Fig. 4B, bottom row). Interestingly, *Patches* also identified genes not explicitly used during annotation. For example, *AW112010*, a long non-coding RNA previously shown to promote the differentiation of inflammatory T cells [49], achieved the highest specificity score for T cells (Fig. 4B, bottom row). This highlights *Patches*’ ability to reveal biologically meaningful signals beyond known markers.

**Figure 4:**
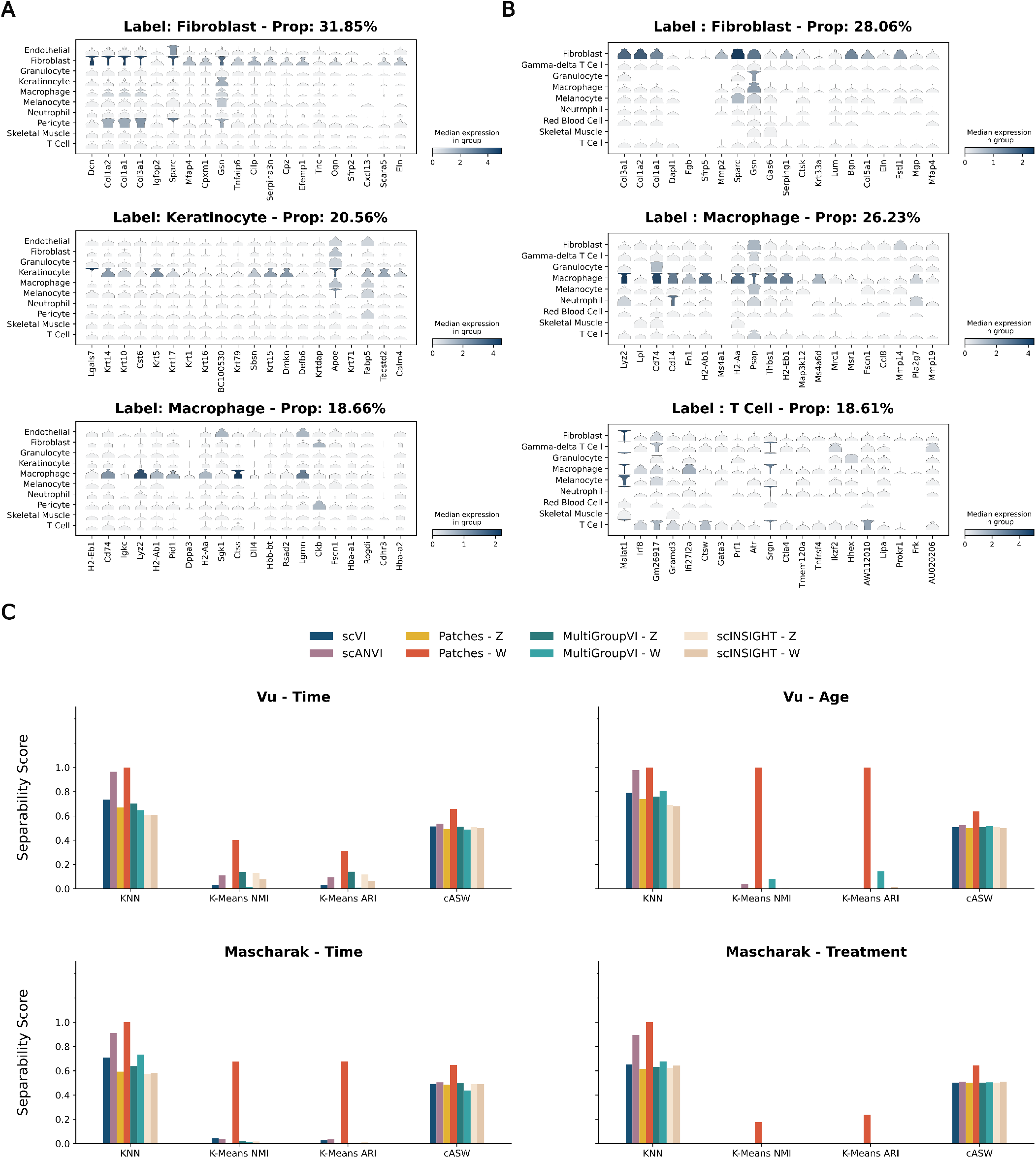
Quantification of the attribute-specific representations of *Patches*. *Patches* can distinguish attribute-specific effects in real datasets. **A, B**, Top 20 marker genes associated with the provided cell type label annotations for the 3 major cell types in the Vu (**A**) and Mascharak (**B**) datasets, obtained from the linear decoder of *Patches*. Violin plots show gene expression distributions across all annotated cell types. **C**, Separability scores for the latent representations obtained by running scVI, scANVI, MultiGroupVI, scINSIGHT and *Patches* on the Vu (top row) and Mascharak (bottom row) datasets. KNN, K-Means NMI, K-means ARI and cASW scores for models were calculated separately for each condition group within each dataset - a higher score indicates higher separability.

Similarly to existing deep learning methods, *Patches* relies on having sufficient examples to learn stable and informative latent structure. To explore how this dependence manifests in practice, we examined performance on the three least abundant cell types (Sup. Fig. S3). As the representation becomes constrained by fewer observations, the interpretability module provides correspondingly less informative summaries. This suggests that, when the goal is to interpret rare cell populations, explicitly subsetting or rebalancing the data may be necessary to obtain reliable representations.

#### Attribute-specific representations show clear separation and interpretability

To assess the ability of *Patches* to separate conditional clusters, we compared its learned latent representations with those generated by four widely used generative models: scVI [16], scANVI [25], MultiGroupVI [50] and scINSIGHT [51]. For each condition group across both datasets, we calculated separability scores based on the encoded observations within the latent spaces of each model (Fig. 4C). We considered four metrics: K-nearest neighbor classification accuracy (KNN), normalized conditional Average Silhouette Width (cASW), K-Means Adjusted Rand Index (K-Means ARI), and K-Means Normalized Mutual Information (K-Means NMI) [52, 44] (*Methods*). A higher KNN score indicates that points with the same attribute are closely positioned in the latent space. Normalized cASW scores reflect cluster compactness and distinctness, with moderate values indicating overlap. K-Means-based metrics convey correspondence between original data attributes and their organization in the latent space. Importantly, as *K* must match the number of attributes for each condition group (*Methods*), achieving a high score requires exactly matching cluster structures between two clusterings.

For cell type annotations, scVI achieved the highest separability scores among the tested models when evaluated using K-Means-based metrics. This is likely because scVI does not incorporate labels during training, instead prioritizing the largest axis of variation when structuring the latent space. As a result, it preserves global cluster structures related to cell types. In contrast, MultiGroupVI, scINSIGHT, and *Patches* are designed to learn structured latent representations. *Patches* in particular organizes variation around combinations of attributes, rather than primarily preserving global structure (Fig. 3, second-to-last column). This results in greater variability across scores for K-Means-based metrics (Fig 4C). However, cells of the same type are still situated closely in the conditional latent space of *Patches*. KNN and cASW scores seem to reflect this, as conditional representations of *Patches* achieve scores similar to scVI (Fig. 4C, right panels). Unlike *Patches*, MultiGroupVI and scINSIGHT are not explicitly designed to isolate cell-type–specific signals into a dedicated latent component. To make this difference explicit, we compared the cell-type–specific representation learned by *Patches* to the common latent components of MultiGroupVI and scINSIGHT (Sup. Fig. S4). Across metrics, this comparison shows that both MultiGroupVI and scINSIGHT tend to encode cell-type–associated gene expression signatures more strongly in their common representations than in their condition-specific components, as reflected by lower separability in *w* relative to *z*. For scINSIGHT, this behavior varies across datasets and metrics: in the aging dataset, separability is comparable between common and condition-specific representations, whereas in the drug treatment dataset, higher separability is observed in the common space. By contrast, scVI, scANVI, *Patches*, and MultiGroupVI exhibit more consistent separability patterns across datasets. These observations highlight a general tension in representation learning for multi-conditional data. Some degree of cell-type signal is expected to reside in shared representations, reflecting gene programs that are conserved across conditions. The goal of explicitly modeling cell-type–specific components, as in *Patches*, is not to eliminate shared structure, but to distinguish signatures that are specific to cell identity from those that are broadly shared across cell types and conditions.

For condition-specific attributes (age, time or drug treatment), scVI showed lower separability scores, comparable to the values observed for the common representations of *Patches* (Fig. 4C, left and middle panels). Additionally, common representations always attain the lowest separability scores across comparisons, highlighting the desired mixing behavior for attributes (Fig. 3, second column). For scANVI, MultiGroupVI, and scINSIGHT, KNN and cASW scores indicate that condition-specific labels are positioned close together in the latent space, mirroring the behavior of the conditional representations learned by *Patches* (Fig. 4C, left and middle panels). This is consistent with the design of these methods, which are provided with condition labels and encouraged to organize their latent spaces to separate combinations of attributes. However, both MultiGroupVI and scINSIGHT exhibit comparable separability in their shared and condition-specific representations, suggesting that age- and condition-related information is not cleanly isolated and may be distributed across components. By contrast, constraining how age- and condition-related information is expressed allows *Patches* to separate these effects more cleanly, while preserving a shared representation that aligns cells across ages and treatments.

Overall, *Patches* effectively models variation across different data attributes. Owing to the additional structuring of its latent space [38] (*Methods*), Patches achieved better separation of latent clusters, leading to higher separability scores.

### *Patches* achieves competitive reconstruction and transfer accuracy in scRNA-seq data

We next assessed the ability of *Patches* to disentangle conditional information while generating realistic gene expression profiles through its decoder. In addition to reconstruction tasks, we tested the model’s capacity to ‘transfer’ gene expression patterns across timepoints, a key challenge in datasets with dynamic conditions.

We split the datasets into training and test subsets and evaluated the reconstruction accuracy across all cells and the three major cell types by proportion. *Patches* achieved reconstruction scores comparable to scVI, which does not incorporate conditional labels, and outperformed scANVI across all comparisons (Fig. 5A). Notably, in the Vu dataset, scANVI performed relatively well for fibroblasts, likely due to their dominance as the majority cell type.

**Figure 5:**
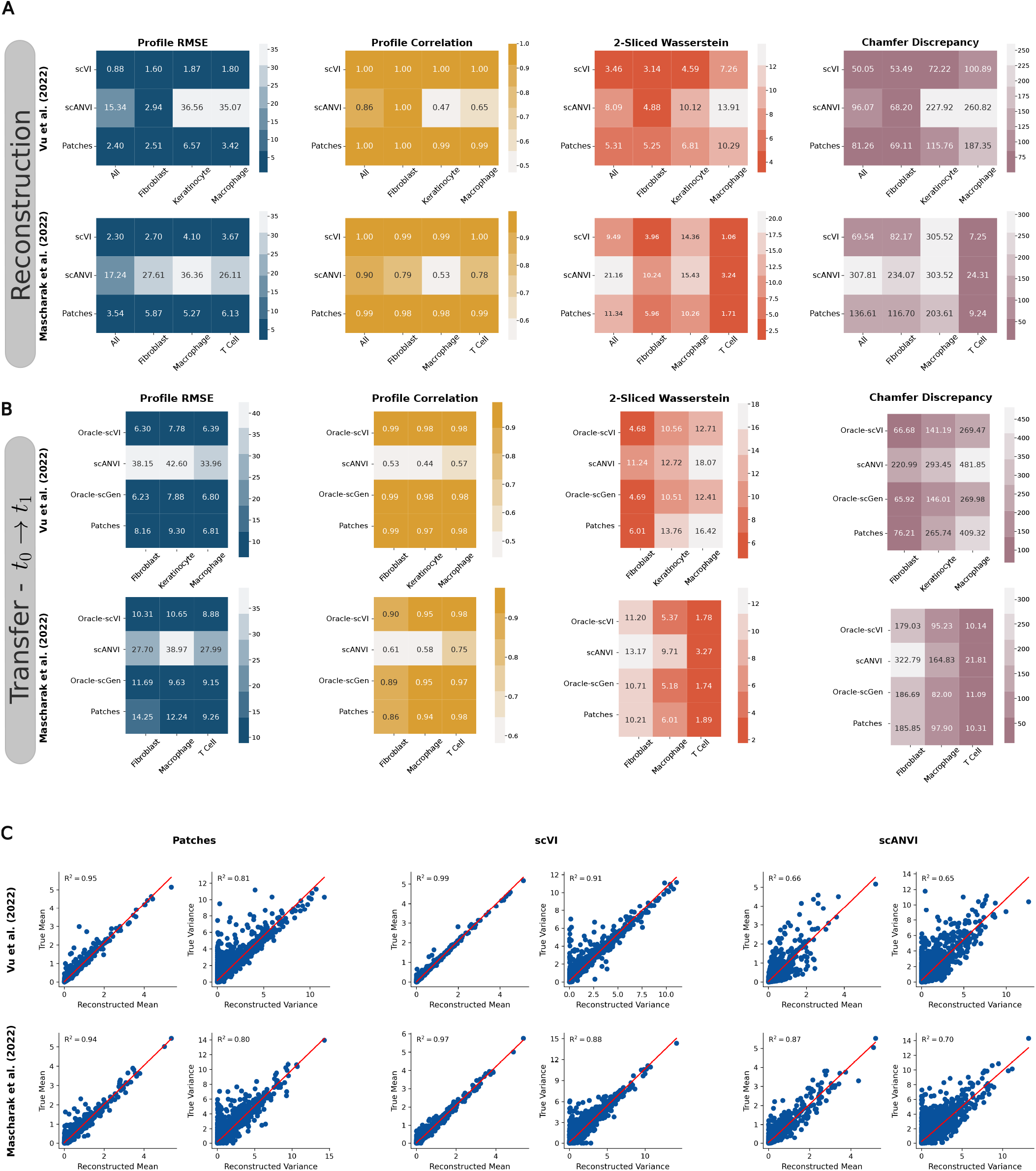
*Patches* performs competitively in generative tasks. Comparison of generative accuracy metrics (RMSE, Correlation, 2-SWD and CD) for scVI, scANVI, scGen and *Patches*. Darker colors indicate better performance. **A**, Reconstruction accuracy on held out cells for the benchmarked models for all cells and the three major cell types for the Vu (top row) and Mascharak (bottom row) datasets. **B**, Transfer accuracy for the benchmarked approaches calculated after transferring held out cells of the 3 major cell types from the first (0DPW for Vu, POD7 for Mascharak) to the subsequent timepoint (4DPW for Vu, POD14 for Mascharak) for the Vu (top row) and Mascharak (bottom row) datasets. **C**, Comparison of *Patches* (left), scVI (middle) and scANVI (right) in recapitulating the true and reconstructed log-mean and log-variance for each gene across the Vu (top row) and Mascharak (bottom row) datasets.

We further tested the model’s ability to ‘transfer’ gene expression patterns across consecutive timepoints (Sup. Fig. S5, Methods). Starting with observations from the first timepoints (0 DPW for Vu, POD7 for Mascharak), we used Patches to transfer these expression profiles to the second sets of timepoints (4 DPW for Vu, POD14 for Mascharak). As proportions of cell types are different across groups, we performed the transfers per cell type for the three major cell types in each dataset. The objective was to understand how these observations might appear if collected at the second timepoint. While Patches and scANVI provide internal mechanisms to facilitate transfer between conditions, scVI was not originally designed for this purpose. To address this, we introduced an augmented version of scVI with oracle information (Oracle-scVI, *Methods*). We also include a similarly augmented version of scGen [29], which was originally proposed for related transfer tasks (Oracle-scGen, *Benchmarks*). In this setup, scVI and scGen were provided with the statistical distance between the source (gene expression from the first timepoint) and the target (gene expression from the subsequent timepoint) in latent space. Because these models do not ordinarily have access to how the mean gene expression changes across unobserved conditions, we call and treat these variants as Oracle models, intended to provide an upper bound on performance for this task. Comparing the four approaches, *Patches* implemented transfer through label change with high accuracy, achieving performance comparable to Oracle-scVI and Oracle-scGen (Fig. 5B) In contrast, scANVI performed poorly across all cell types, with its best performance observed for T cells in the Mascharak dataset.

To quantify how well the models reproduced biological variation, we compared the mean and variance of reconstructed gene log-counts to the original log-counts. Consistent with the reconstruction accuracy results, Patches and scVI effectively recapitulated gene-level variability, outperforming scANVI (Fig. 5C). The weaker performance of scANVI may stem from its reliance on a single one-hot encoded label to represent multiple condition groups, a strategy that does not scale effectively for datasets with many conditions which might combinatorially overlap.

### *Patches* identifies key genes and pathways driving tissue repair and aging dynamics

Next, we explored how the latent representations learned by *Patches* contribute to separating cells according to their attributes. To this end, we first extracted the first PC from both common and conditional latent representations across the wound healing datasets considered. Then, for each gene, we computed alignment scores (*R*^2^) by correlating the obtained PC values with the original transcript counts, which are shown on the Y-axis in Figure 6, Panel A. Simultaneously, we performed differential gene expression tests using the normalized count matrix, grouping observations by condition-specific attributes (e.g., age and time for the Vu dataset [4]; treatment and time for the Mascharak [48]). Corresponding results from standard differential gene expression analyses of the preprocessed dataset are provided in Supplementary Figures S6 (age and time) and S7 (treatment and time). The resulting significance scores (−*log*10 adjusted P-values) are plotted on the X-axis. As expected, the common latent representations do not encode condition-specific information, reflected by the lower correlation between alignment scores and differential expression significance, as well as the low separability score (quantified using the Average Silhouette Width). Furthermore, the alignment score (Fig. 6A) indicates that the common representations *z* account for a substantial portion of the variation remaining after conditioning on *w*. The relative magnitude of this component is consistent with the view that condition-specific effects are often smaller than shared transcriptional structure in real experimental settings.

**Figure 6:**
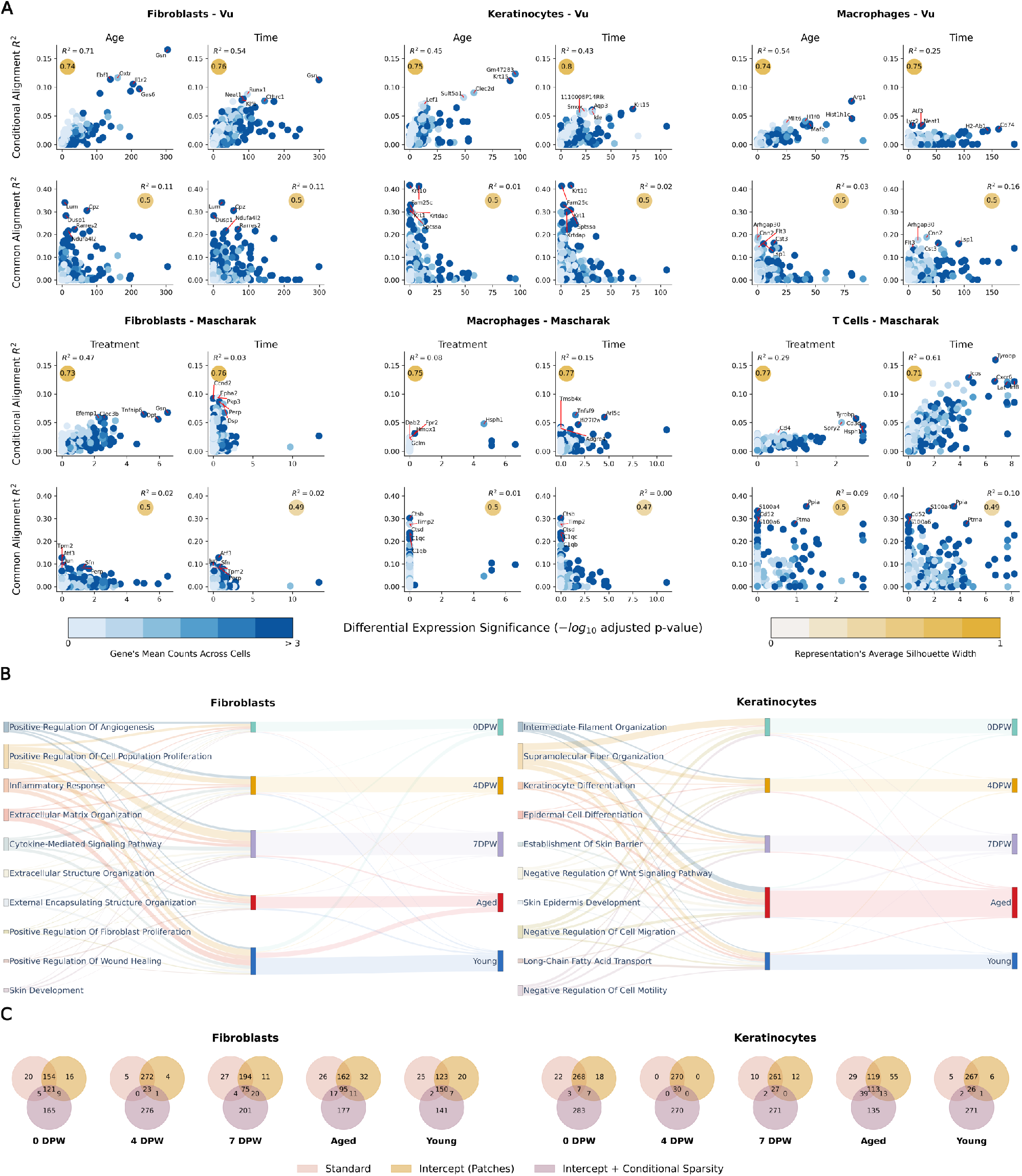
Extracting condition-specific interpretations with *Patches*. **A**, Alignment *R*^2^ values from conditional and common representations (Y-axis) plotted against differential gene expression significances obtained by comparing expression of genes across condition labels (X-axis) for the 3 major cell types (columns) of the Vu (top row) and Mascharak (bottom row) datasets. Yellow circles indicate separability score for the embedding obtained by *Patches* (Average Silhouette Width). We annotate the top 5 genes attaining the highest alignment *R*^2^. **B**, Sankey diagram of the top 10 significantly enriched GOs for the top 10% (*n* = 300) genes in fibroblasts (left) and keratinocytes (right) of the Vu dataset. Genes appearing in the top 10% across multiple attributes were labeled with the top scoring attribute, but shown connected to other attributes through links. Link bandwiths between GOs and attributes represent the flow of attribute scores. **C**, Venn diagrams of top 10% genes for each attribute in the Vu dataset by applying linear decoders with different statistical architectural choices (intercept, sparsity) for fibroblasts (left) and keratinocytes (right).

The scores derived from the conditional latent representation can be interpreted in two ways. First, they may reflect a direct influence of the condition on gene expression. Specifically, if a condition-specific factor (e.g., age or treatment) directly impacts gene expression, this would result in higher *R*^2^ values in the linear fit of the alignment-differential expression plots. This indicates that variations in gene expression align closely with condition-specific latent changes, as observed for aging fibroblasts in the Vu dataset. Second, the scores can be indicative of a nonlinear or indirect relationships. Namely, in cases where values are lower, the condition-specific changes are not directly captured by differential expression analysis. This suggests a more complex, potentially nonlinear relationship between latent representations and expression values.

We found several genes with high alignment scores that were associated with wound healing and aging phenotypes in a cell type-specific manner. In fibroblasts, key genes such as *Gsn, Oxtr, Ebf1, Il1r2*, and *Gas6* exhibited strong associations with processes central to tissue repair and regeneration. *Gsn* plays a critical role in actin cytoskeletal remodeling, enabling fibroblast migration during wound healing [53]. *Oxtr* is involved in tissue regeneration and homeostasis [54], while *Ebf1* regulates cell differentiation [55], and *Gas6* promotes fibroblast survival and tissue repair via TAM receptor signaling, with a notable role in age-dependent fibrosis [56].

Similarly, aging effects were prominent in keratinocyte populations. Here, the highest-scoring altered genes included *Krt15, Lef1, Clec2d, Sult5a1*, and *Gm47283. Krt15*, a marker of hair follicle stem cells [57], and *Lef1*, a regulator of the Wnt pathway, suggest impairments in follicular stem cell maintenance and regenerative capacity [58, 59]. *Clec2d* reflects immune modulation and aging dependent inflammation [60], while *Sult5a1* indicates altered metabolic and hormonal processing alongside aging in the interfollicular epidermis (IFE) [61, 62]. The uncharacterized *Gm47283* may implicate RNA-based regulatory mechanisms. Together, these findings demonstrate *Patches*’ ability to identify genes linked to critical pathways involved in tissue repair, stem cell exhaustion, immune dysregulation, and age-related metabolic changes across fibroblast and keratinocyte populations.

Finally, we incorporated a linear decoder into *Patches* to derive interpretable contributions of individual genes to the latent representations [19]. Using this computational framework, we analyzed the two major cell types in the Vu dataset — fibroblasts and keratinocytes. Group-conditional scores were computed for each gene, capturing linear relationships between condition-specific attributes (e.g., cell type, age, time) and gene expression levels (*Methods*). Genes were ranked by attribute and the top 10% of genes were selected for downstream analyses, including gene ontology (GO) enrichment analyses [63] (Fig. 6). We found gene programs associated with ECM organization were more enriched in fibroblasts rather than in keratinocytes in the aged groups, corroborating previous findings regarding ECM dysregulation in ageing fibroblasts [64]. Conversely, gene programs specific to keratinocytes showed a larger contribution to intermediate filament organization programs, further supporting the fact that aging significantly impacts intermediate filament organization during tissue repair [65, 66].

Using a linear decoder and a new in-house dataset [67] of unwounded and DPW7 mice (*Datasets*), we assessed the performance of the interpretability module of *Patches* in a real experimental setting. To this end, we carried out an analysis analogous to the identifiable simulation study (*Methods*). After fitting *Patches* with a linear decoder on the full dataset, we quantified gene-level time-specific effects by computing the mean absolute deviation (sAD) of time-associated scores across genes for the unwounded and DPW7 conditions (Sup. Tab. 3, 4). We then ranked the upregulated genes by decreasing absolute deviation. GO enrichment results for top 10% of ranked genes are presented in Supplementary Figure S7. Here *Patches* successfully identifies signals related to wounding and ECM degredation across cell types (Sup. Fig. S8A). Top terms in Supplementary Figure S8B,C denote possible cell-cell communication mechanisms, which are known to be important for wound healing [5]. Further ECM association is observed in Supplementary Figure S8D. We validate the overlap of certain genes with respect to time after injury among the top 10%, such as *Ebf1* and *Col1A2*. Some top genes among the 10% include: members of the various Interleukin families such as *Il18r1, Il34, Il1a, Il20ra*; additional members of the Collagen family such as *Col1a1, Col25a1, Col3a1, Col12a1, Col5a2, Col4a6, Col5a1*; members of the Keratin family such as *Krt5, Krt79, Krtap19-4, Krt14, Krtap4-2, Krt24*; and finally chemokines such as *Cxcl2, Cxcl3, Ccl3, Ccl4*.

To assess the impact of modeling choices on gene-specific interpretability scores, we examined three variants of the linear decoder making up *Patches*’ interpretability module. In addition to a standard linear decoder architecture [19], we also considered an extension with an explicit intercept term, and a model incorporating both an intercept and an ℒ_1_ sparsity penalty on the conditional coefficients (*Methods*; *Interpretability module*). We compared the top 10% of genes from the ranked lists generated by each implementation.

Across the three decoder variants, Patches consistently identified top-scoring genes in a cell-type and age-specific manner, highlighting their biological relevance to wound healing and aging processes (Fig. 6C). In fibroblasts, the intersection of top genes for aged samples included *Gzmc, Scube3, Cilp, Smoc1*, and *Apoe*. In particular, *Scube3* has been linked to ECM organization and growth factor signaling [68], and both *Cilp* and *Smoc1* are ECM-associated proteins, with *Cilp* implicated in age-related fibrosis [69, 70] and *Smoc1* promoting fibroblast migration and repair [71]. Notably, *Apoe* is a key regulator of lipid metabolism and inflammation, both of which are altered in aging fibroblasts [72]. In young fibroblasts, top genes included *Gas6, Gsn, Il1r2, Hbb-bt*, and *Cilp*. Among them, *Hbb-bt*, a hemoglobin beta subunit, may reflect metabolic activity associated with fibroblast responses [73]. The consistent appearance of *Cilp* across both age groups underscores its importance in ECM regulation and tissue remodeling.

In keratinocytes, *Fabp5, Krt25, Krt35, Tchh*, and *Slc30a1* were identified as top genes for aged samples across algorithmic variants. Notably, *Krt25* and *Krt35* are keratin genes associated with cytoskeletal stability, which are reflecting age-related changes in keratinocyte structure and function [74, 75], while *Tchh* (trichohyalin) is involved in keratinocyte differentiation and hair cell maturation [76, 77, 78, 79], and *Slc30a1* encodes a zinc transporter linked to cellular stress responses and proliferation [80]. For young mice, Patches identified *Slc27a1, Tgfa, Col1a2, Terf2ip*, and *Calml3. Slc27a1*, a fatty acid transporter, and *Tgfa*, a growth factor involved in epidermal proliferation, are involved in active lipid metabolism and regeneration processes in younger cells [81, 82]. Similarly, *Col1a2*, a collagen gene, underscores ECM production and tissue integrity [83], while *Calml3* is linked to calcium signaling, which plays a role in keratinocyte differentiation and proliferation [84].

Computationally, adding the intercept term to the standard approach resulted in minimal changes to the ranked gene lists. However, introducing the sparsity term in the loss function altered the lists to varying degrees, depending on the condition-specific attributes (Fig. 6C). This observation suggests that many genes may contribute nonlinearly to the attribute-specific representations. Alternatively, it highlights the need to carefully document and scrutinize non-trivial modeling choices, as such choices can strongly influence the interpretability and reproducibility of generative models applied to scRNA-seq data [85].

## Discussion

Here we introduce *Patches*, a generative deep learning framework designed to learn both shared and condition-specific gene expression representations from experiments with observations collected across multiple conditions. In the context of wound healing, this approach provides a powerful tool for understanding the impact of cell type, age, or treatment. Specifically, *Patches* enables both qualitative and quantitative analyses of single-cell datasets in complex experimental designs involving multiple conditional labels, providing researchers with deeper insights into the cellular dynamics of wound healing. *Patches* leads to visualizations that separate gene expression signals into common and distinct nonlinear patterns, complementing existing dimensionality reduction techniques. This separation is achieved through information-theoretic priors, which promote disentangled representations. Furthermore, *Patches* can handle data collected from experiments involving combinatorial condition groups, where not all the cells are observed across all the groups. Finally, its interpretable design facilitates an understanding of how individual genes contribute to the learned data representations. Using Patches, we identified several unique age-related changes, including notable alterations in cellular pathways, extracellular matrix components, and gene expression patterns. Among these, a novel observation was the alteration of *Apoe*, a key regulator of lipid metabolism and inflammation, in aging fibroblasts. This finding may provide a mechanistic explanation for the altered lipid homeostasis, increased inflammatory signaling, and functional decline observed in aged fibroblast populations [86, 4, 87].

Despite its strengths, *Patches* presents computational challenges, including longer training times compared to simpler models like scVI. This is partly due to the use of adversarial discriminators and the additional training epochs required to ensure disentangled latent representations. Furthermore, while the linear decoder enables interpretability, its expressivity may limit performance in complex generative tasks such as cross-condition imputation. Nonetheless, this trade-off is offset by its ability to generate biologically interpretable insights. The form of disentanglement pursued by *Patches* reflects the inherent difficulty of partitioning variation into label-associated effects, captured by *w*, and remaining structure, captured by *z* [88, 89]. *Patches* achieves this through heuristics, such as using an adversarial classifier and conditional bottlenecks on the dimensions of *w*. As a result, model behavior depends on hyperparameter choices (*Methods*) and sufficient sample size, a dependence illustrated in Supplementary Figure S3.

A promising future application for *Patches* is its application to non-healing wounds and cross-species comparison of wound healing dynamics. Direct studies of wound healing in humans are limited by ethical and practical constraints, yet substantial single-cell transcriptomic datasets from unwounded and diseased human tissues are readily available [9, 90, 91]. *Patches* could bridge this gap by leveraging transcriptomic patterns from unwounded human samples and integrating data from animal models, where wounding can be experimentally induced. This approach could provide critical insights into the conserved and divergent mechanisms underlying wound healing across species. Such cross-species predictive analyses hold significant potential for improving the translation of findings from model organisms to human biology, paving the way for the development of targeted and more effective therapeutic strategies for wound healing. Further extending Patches to accommodate additional data modalities, such as spatial transcriptomics or chromatin accessibility data could enhance its utility in studying dynamic biological processes. Specifically, spatially resolved data will allow us to couple a cell’s identity and its gene expression programs with the context of its neighbors, thereby revealing the cell–cell interactions that drive coordinated behavior during wound healing. Lastly, improving scalability and computational efficiency would enable broader adoption in large-scale studies.

The ability to disentangle shared and condition-specific transcriptomic patterns makes *Patches* a valuable tool for investigating multi-condition biological systems. Its application extends beyond wound healing, with potential use cases in developmental biology, aging research, and response to therapeutic interventions. As datasets become increasingly complex and multifaceted, *Patches* provides a critical framework for extracting interpretable insights, driving discoveries in systems biology and translational research.

## Methods

### Overview of *Patches*

*Patches* is a representation learning framework which extends traditional VAE models in various important ways. First, *Patches* incorporates a hierarchical latent space to improve visualization and generative accuracy. Moreover, the underlying latent variables are organized into common and conditional components. The conditional component captures variation associated with the provided labels, whereas the common component accounts for the remaining structure in the data. Information theoretical constraints ensure the separation between common and condition-specific latent representations. Finally, *Patches* allows interpretability through incorporating a linear decoder [19] to identify gene expression signatures driving both common and condition-specific cell states. In detail, our framework integrates and extends several key concepts from computational biology and machine learning into a unified approach:

#### Deep Generative Models

Deep generative models, such as scVI [16] and its extensions [25, 31, 27, 28], have demonstrated success in generating highly realistic gene expression profiles.

#### Latent Space Structuring

Organizing the latent space can significantly enhance model performance. Hierarchical structuring of the latent space improves visualizations [23], while conditioning the latent space on discrete categories (e.g., chemical interventions or cell types) enables learning gene expression patterns specific to attributes in the data[25, 34, 92, 30].

#### Challenges in Disentangled Representations

Learning conditional disentangled latent representations is inherently challenging [38, 39]. In some cases, it may even be theoretically unattainable [93]. Most importantly, organizing the latent space only as a mixture of conditions can overlook important biological factors. For instance, in wound healing studies, gene expression signatures may be present irrespective of whether the cells originate from old or young mice, alongside age-specific signatures. Previous work has shown the benefits of incorporating both common and conditional latent representations in generative [33] and non-generative models [35]. However, these models necessitate explicit cross-modality alignment, which is impractical when observing complex scenarios like wound healing in both young and old mice simultaneously.

#### Regularization for Disentanglement

To ensure the disentanglement and the identifiability of common and conditional latent representations, additional regularization is necessary. This can be achieved through imposing various constraints [38, 39, 94].

#### Improving Interpretability

End-to-end deep learning models often suffer from poor interpretability. This can be enhanced by incorporating linear decoding layers [19].

### Generative model

#### Notation and setup

In the following, *n* indexes the *N* cells and *g* indexes the *G* genes in a given study. The corresponding gene expression values are denoted by *x*_*ng*_ for each cell *n* and gene *g*. We encode observation labels through a two-level hierarchy consisting of *C* condition groups each with *S*_*i*_ attributes, where *i* = 1, 2, … *C* indexes the *C* classes. For example, ‘age’ represents a class or a condition group, and ‘young’ or ‘old’ represent its *S*_*age*_ = 2 attributes. The total number of attributes is denoted by *K*, where *K* = *S*_1_ + *S*_2_ + … + *S*_*C*_, and each attributes is indexed by *k*. The label information about the observation *n* is encoded in a binary vector **y**_*n*_ ∈ {0, 1}^*K*^. This binary vector is a concatenation of *C* one-hot encodings, unit vectors 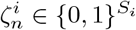, one for each class *i* = 1, 2, … *C*.

As an example, consider observations collected from a dataset with two annotated cell types (fibroblasts and keratinocytes) across three different timepoints (*t*_1_, *t*_2_, *t*_3_) and from mice of different ages (young and old). This sample can be represented with a binary encoding consisting of three condition groups: cell type (*S*_*Cell Type*_ = 2), timepoint (*S*_*Time*_ = 3) and age (*S*_*Age*_ = 2). In this case, let **y**_*n*_ = [1, 0, 1, 0, 0, 1, 0] represent a fibroblast collected from an uninjured, young mouse. This is concatenation of three unit vectors 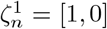 for the cell type, 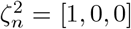 for the timepoint and 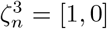 for the age. Note that a sample with the label encoding **y**_*n*_ = [1, 0, 1, 1, 0, 1, 0] would be inadmissible since it would imply that the data comes from two timepoints simultaneously.

#### Model

Each gene expression count *x*_*ng*_ is generated using a hierarchy of latent variables informed by the label encodings **y**_*n*_ under which the cell *n* was collected, as follows:

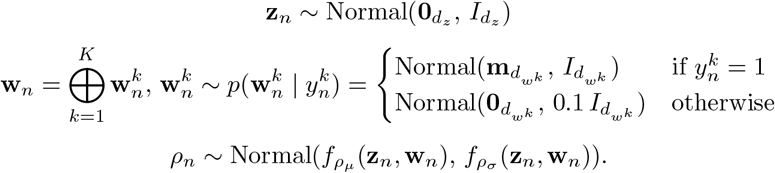

Here, **m** is a constant vector of size 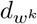 that controls the separation between common and conditional latent variables. The latent variable **z**_*n*_ has a standard multivariate normal prior and reflects the portion of the cell’s identity that is independent of its conditional labels 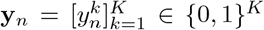. On the other hand, each 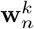 is drawn from one of two multivariate normal distributions, depending on whether the 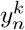 governing the corresponding 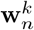 is 1 or 0. Intuitively, this provides a mechanism to turn the corresponding conditional subspace 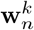 ‘on’ or ‘off’ through the value of 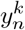. Then, for a cell *n*, **w**_*n*_ is defined as the concatenation (⊕) of all such 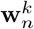. The functions 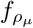 and 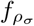 are used to generate a cell’s latent representation *ρ*_*n*_ using its conditional latent representation **w**_*n*_ and common latent representation **z**_*n*_. The final gene expression profiles are generated following methods commonly used in latent variable models for single-cell RNA-seq data [42, 16, 25]:

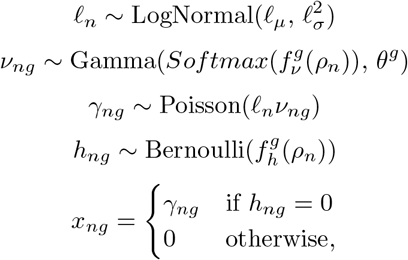

where *ℓ*_*n*_ are cell-specific scaling factors centered around the observed log-library size and 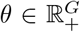 are gene-wise inverse dispersions. The layers *f*_*ν*_ and *f*_*h*_ map the latent representations back into the gene space while accounting for dropouts [42, 16, 25]. The resulting distribution of the gene counts is ZINB, where the zero-inflation probabilities for each gene are encoded by *f*_*h*_.

### Inference and optimization

The posterior distribution of the model latent variable given gene expression profiles 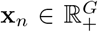 and their corresponding sample attributes **y**_*n*_, that is *p*_*ϕ*_(**w**_*n*_, **z**_*n*_, *ρ*_*n*_, *ℓ*_*n*_ | **x**_*n*_, **y**_*n*_), is intractable [16, 25, 95]. We compute an approximation via stochastic variational inference [16]. In detail, the resulting variational distribution *q*_*ϕ*_(**w**_*n*_, **z**_*n*_, *ρ*_*n*_, *ℓ*_*n*_ | **x**_*n*_, **y**_*n*_) is factorized into neural networks with parameters *ϕ*:

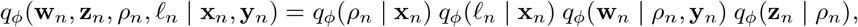

where, for simplicity, we drop the batch indicators *s*_*n*_. Similarly to [16], *p*(*x*_*ng*_ | *ρ*_*n*_, *ℓ*_*n*_) has a closed form density, following a ZINB distribution. Hence, inference over the random variables *ν*_*ng*_, *h*_*ng*_ and *γ*_*ng*_ is not needed. Applying Jensen’s inequality to the approximate posterior, we obtain the variational lower bound which constitutes the first part of our objective:

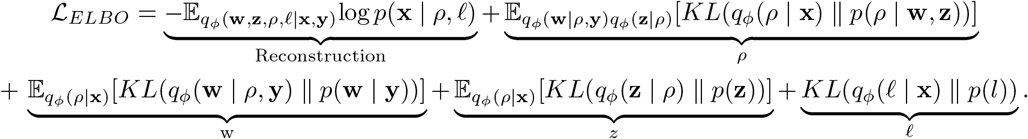

This variational objective encodes the assumption that the common latent representations **z** are independent of the conditional labels **y**. However, in practice, minimizing this objective does not necessarily lead to solutions that satisfy the independence condition [38]. Additional constraints are necessary to ensure that all information correlated with **y** is removed from **z**. To do so, we employ an information-theoretic strategy and introduce a condition-dependent discriminator loss:

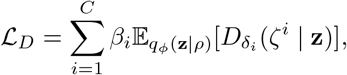

where the parameters *β*_*i*_ weigh the importance of individual conditions and *δ*_*i*_ are the parameters of the neural networks corresponding to the discriminators of each condition *ζ*^*i*^. During training, the discriminators try to maximize their accuracy of predicting the correct attribute for the corresponding condition class (e.g, predicting ‘early’ for the condition class ‘healing stage’) only from the common latent representations **z** of the given sample:

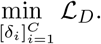

Minimizing this objective would result in representations **z** that are suboptimal: They would retain as much information about the conditional labels **y** as possible, making them difficult to distinguish from condition-specific representations **w**. Since we seek the opposite, our final objective actively penalizes the representations that are favorable for the discriminator loss:

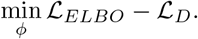

### Cross-condition transfer

*Patches* is designed to allow switching between any condition by treating each condition group as a separate entity in the label encoding **y**_*n*_. Cross-condition transfer for a cell **x**_*n*_ with target conditional label **y**_*t*_ using a trained instance of *Patches* is performed in three steps:

1. Encode **x**_*n*_ into the shared latent space **z**_*n*_.
2. For the conditional identity, either (1) sample a **w**_*t*_ using **y**_*t*_, or (2) construct any **w**_*t*_ of choice [38].
3. Decode the transferred gene expression using the concatenated representation [**z**_*n*_, **w**_*t*_].

Step 2 provides substantial flexibility in defining **w**_*t*_, in that any custom 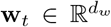 can be provided during transfer. Alternatively, multiple **w**_*t*_ instances can be sampled to explore a range of possible generation outcomes [38]. For our experiments, however, we sample a single **w**_*t*_ using **y**_*t*_ to minimize time spent with computations.

### Interpretability module

*Patches* can optionally use a linear decoder for the generative process, allowing it to associate the expression pattern of each gene *g* with the condition attributes 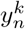 [19]. In this mode, the intermediate variable *ρ* is excluded, and the function 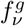 is linear. This option is not suitable for cross-condition prediction, as the linear decoder’s limited expressivity may hinder accurate gene expression generation.

As before, let 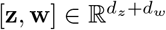 be the concatenation of latent representations, *α* ∈ ℝ^*g*^ be the optional intercept term (which does not appear in the standard linear decoder implementation in [19]), and 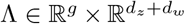 be the matrix linking gene expression profiles to their latent contributions. To encourage interpretability, the generative model can be rewritten as:

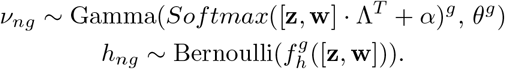

By construction, Λ is a concatenation of two sub-matrices 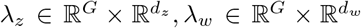 along the columns, which correspond to the common and conditional weights per gene specifically. Additionally, *λ*_*w*_ itself is a concatenation of *K* sub-matrices of the form 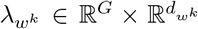 along the columns. The conditional sparsity penalty is an ℒ_1_ loss on the conditional coefficients *λ*_*w*_ defined as:

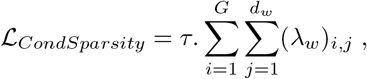

where we chose *τ* = 1*e* −4 across all experiments. To have a single continuous value per gene that serves as an attribute-specific score, we report the sum of weights across the attribute dimensions 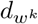.

### Architecture and hyperparameters

#### Architecture

*Patches* uses neural networks to parameterize many of its components: all the encoders used for the approximate variational posterior (*q*(**w, z**, *ρ, ℓ* | **x, y**)), the generative functions 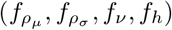, and the discriminators 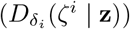. In detail:

- The encoders for the approximate variatonal posterior and the generative functions are neural networks with *n*_*hidden*_ = 2 hidden layers. Each hidden layer is a fully-connected layer of *d*_*hidden*_ = 128 neurons, followed by a *BatchNorm1d* layer of the same size and then a *ReLU* activation.
  The inputs for the encoders are determined by the variables being conditioned on (*Inference and optimization*). The outputs, similarly, correspond to the parameters for the outcome variable distributions. If the parameters are required to be non-negative, the outputs are passed through an additional *Softplus* layer.
  The input for 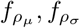 is the concatenation of latent representations **z** ⊕ **w** in this order. The input for *f*_*ν*_, *f*_*h*_ is the latent representation of the cell identity *ρ*. The outputs of 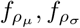 are respectively the mean and variance for the reparameterized cell identity *ρ*. The outputs of *f*_*ν*_ are passed through an additional *Softmax* layer (as in [16]) to encode the mean gene expression proportions irrespective of total counts. The outputs of *f*_*h*_ represents the log-odds for the probability of a technical dropout event for each gene.
  For all neural networks, the inputs are passed through hidden layers, and then a final fully-connected layer.
- The discriminators 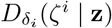 are neural networks formulated in the same fashion with the same hyperparameters. The input representations **z** are passed through the hidden layers, and then a final fully-connected layer. The discriminator output represents log-odds for the probability of belonging to the condition *ζ*^*i*^.

#### Training procedure

During training, the model receives as input the gene expression 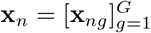 of a cell *n*, along with its label encoding 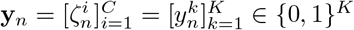. These inputs are passed through an encoder *q*(*ρ*_*n*_, *l*_*n*_ | **x**_*n*_), which parameterizes cell identity *ρ*_*n*_, independent of library size, and a value *ℓ*_*n*_ corresponding to the library size of the cell. The cell identity *ρ*_*n*_ is then decoupled into two components: a common representation **z**_*n*_, which is agnostic to the encoding 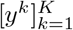, and a conditional representation 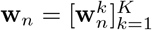. To enforce the independence between **z**_*n*_ and **y**_*n*_, discriminators 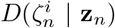 attempt to predict the cell’s conditions 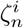 based on its latent representation **z**_*n*_. The latent representations are further used by the decoders 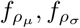 to generate reconstructed cell profiles 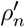, which are then mapped back to the original gene expression space by the decoders *f*_*ν*_, *f*_*h*_ and the encoded library size *ℓ*_*n*_.

#### Implementation details and hyperparameters

*Patches* was implemented using the Pyro probabilistic programming framework [96]. For all real datasets, we used latent dimensions of *d*_*z*_ = 10 for condition-agnostic representations *z*, and 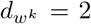 for each attribute-specific representation **w**^*k*^. The total latent dimension was set as *d*_*ρ*_ = *d*_*z*_ + *d*_*w*_. In detail, for the Vu dataset [4], this corresponds to attribute-specific representations with dimensions 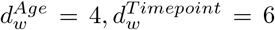 and 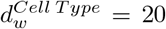. Similarly, for the Mascharak dataset [48] we learned attribute-specific representations with sizes: 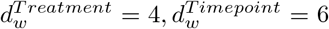 and 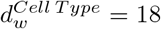. All the fully connected layers of the neural networks were initialized using Kaiming initialization [97], while the *BatchNorm* layers followed PyTorch default settings with scaling initialized to a vector of **1** and biases to a vector of **0** [98]. The multivariate normal distributions of each **w**^*k*^ were parameterized by a constant vector **m** set to **3**, in line with [38]. To ensure consistent scaling, we normalized the loss function by the number of genes *g* and the mini-batch size *d*_*b*_. Optimization was performed using the Adam optimizer with parameters *γ* = 0.001, *ϵ* = 0.01 and *β*_1_, *β*_2_ = [0.9, 0.999]. All models were trained for 2000 epochs on the Vu dataset, and for 20000 epochs on the Mascharak dataset.

#### Details on the choice of latent dimensions

*Patches* includes an adversarial component to ensure the common representation *z* is uninformative of conditions. However, this alone does not prevent common information from leaking into conditional representations *w*. In the degenerate case 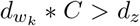 where the number of condition groups (eg. age, treatment, days post wounding) and corresponding dimensions are large enough, there is nothing preventing the model from modeling the data entirely through *w*. Previous theoretical results provide explanations for this in the linear case [99], from which we adapt similar guidelines. In applications of *Patches*, since the maximum number of informative dimensions (ie. those not sampled near **0** due to the conditional prior) for *w* will be 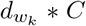. Hence, we ensure that 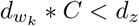 for both our examples, and encourage future applications to do the same.

### Illustrative examples

This section provides a detailed explanation of the methods used to implement the simulations that generated the results presented in Figure 2B-G.

#### Synthetic attribute separation model

We considered the dataset 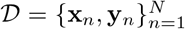 for *N* = 8, 000 where **x**_*n*_ ∈ ℝ^30^ and 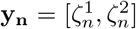 for conditional groups 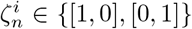. The total number of sampled points was distributed equally across different possible combinations of 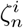. Each data point **x**_*n*_ was constructed through the following procedure:

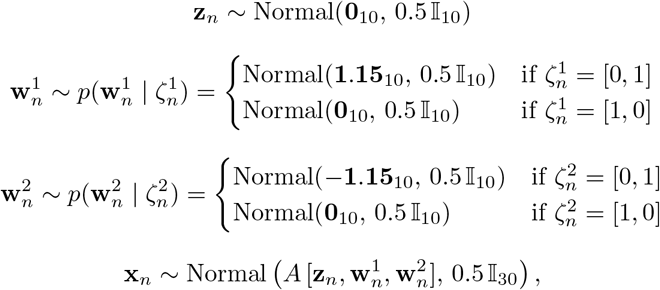

where **b**_10_ denotes a 10-dimensional vector with all entries being equal to the constant value *b*, and 𝕀_10_ is the 10 × 10 identity matrix, *A* is a 30 × 30 identity matrix, and 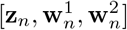 denotes the concatenation of the latent vectors.

#### Swiss roll

We used the previously implemented `make_swiss_roll_dataset` function [44, 45] to generate *N* = 10, 000 points situated on a Swiss roll in three dimensions (*X, Y, Z*). Following point generation, we sliced the roll from the midpoint along the *Y*-axis – at *Y* = 10 – and assigned labels to points to reflect their slice of origin. This resulted in a dataset with a single condition group having two attributes. 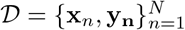 where **x**_**n**_ ∈ ℝ^3^ and **y**_**n**_ = *ζ*_*n*_ ∈ {[1, 0], [0, 1]}.

#### Identifiable simulation study

We rely on an identifiable simulation framework introduced by Wang et al. [46], in which `Splatter` [47] is used to generate data with controlled structure. In this setup, cell–cell similarity is governed by two parameters varied over a grid: *t*, which modulates similarity across cell types, and *s*, which modulates similarity across experimental conditions. Additional implementation details are provided in the original work under the *Simulation setup* section [46]. We consider three cell types and two conditions (Sup. Fig. S1).

### Datasets

We used publicly available single-cell datasets representing various wound healing contexts. These datasets included samples exposed to distinct temporal, chemical, or pathological conditions. We follow a similar approach for every dataset to preprocess and annotate the raw objects prior to running the models.

#### Vu dataset

The Vu dataset [4] contains data for *N* = 26, 966 cells after preprocessing, with samples collected from 10 seven-week-old and six 88-week-old wild-type female mice. Each sample corresponds to data from single mouse and one of three conditions: unwounded (UW), four days post-wounding (4DPW), or seven days post-wounding (7DPW). We only consider the samples from v3 sequencing runs to limit the influence of chemistry-related batch effects. As per the original analysis conducted, we keep cells with at least 200 and at most 5000 non-zero genes and with less than 15% of gene counts arising from mitochondrial genes. We use the standard `scanpy` analysis pipeline [13, 100], assign broad cell type labels based on the output of the `rank_genes_groups` function, and finally validate the cell type labels assigned by comparing with markers from previous literature.

#### Mascharak dataset

The Mascharak dataset [48] consists of *N* = 2, 284 cells after preprocessing. Samples were obtained from mice treated with a PBS control or with verteporfin at five key timepoints (UW, 2DPW, 7DPW, 14DPW, 30DPW). As UW cells are not treated with any substance, we did not choose to include these cells in the final dataset considered. Since antibody counts for demultiplexing were missing from cells belonging to the 2DPW timepoint in the raw data, we instead only use the cells from 7DPW, 14DPW and 30DPW. For demultiplexing, we used the implementation of `hashsolo` [101] provided in [13]. After matching the sample metadata, we apply the standard `scanpy` analysis pipeline [13] to assign broad cell type labels based on the output of the `rank_genes_groups` function, and validate these cell types by comparing with markers from previous literature.

#### snRNA-seq dataset of unwounded and DPW7 mice

To validate signals of tissue repair captured by *Patches*, we apply it to a new in-house single-nuclei RNA-sequencing (snRNA-Seq) [67], consisting of *N* = 21, 176 cells. Samples were obtained from unwounded and seven days post-wounding (DPW7) mice, which were then preprocessed and annotated through `R` using the `Seurat` package. Using the standard analysis pipeline, we kept cells with at least 200 and at most 5000 non-zero genes and with less than 10% of gene counts arising from mitochondrial genes and less than 50% of gene counts arising from ribosomal genes.

#### Data processing

Gene expression count matrices were analyzed with the `scanpy` [13] Python package (version 1.10.0). Separate samples belonging to each dataset were merged into a single `anndata` [102] object prior to preprocessing. After preprocessing and before running the models, for any group of cells considered, we retain the top *g* = 3, 000 highly variable genes. Specifically, we obtain these genes using the `highly_variable_genes` function provided through `scanpy` with default parameters.

### Benchmarks

*Patches* is uniquely designed to (a) learn both shared, condition-agnostic representations and condition-specific representations, and (b) transfer information across different combinations of conditions. While these capabilities are specific to *Patches*, we extended existing methods with oracle-like features for comparison, allowing us to comprehensively evaluate the performance of our framework.

#### Oracle-scVI

The original scVI framework [16] provides a biologically informative latent space, but is not inherently designed for condition transfer. To enable this, we define an attribute manipulation function [38, 29]. Let **y**_*s*_, **y**_*t*_ represent the source and target conditions, and *µ*_*s*_, *µ*_*t*_ represent the means of latent representations for cells collected under these conditions. Switching a cell **x**_*n*_ from the source to the target condition is defined as:

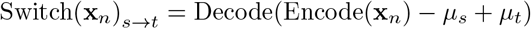

In this process, the cell’s gene expression profile is encoded, transformed via arithmetic in the latent space to enable condition transfer, and decoded to reflect the new condition.

#### scANVI

A semi-supervised version of scVI, scANVI, allows learning latent representations informed by user-specified cell annotations [25]. However, as scANVI is not inherently designed for condition transfer, we adapted it by making the model fully supervised with respect to one hot encodings of attribute combinations 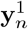. This allows us to define a function for switching between these conditions [38]. In detail, letting 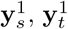 represent the one-hot encoded source and target conditions, switching between conditions for a cell **x**_*n*_ can be expressed as:

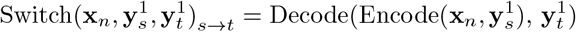

This way, we encode the cell using its source condition and switch the label to the target condition during decoding.

#### Oracle-scGen

scGen [29] extends scVI [16] by incorporating latent arithmetic for predicting counterfactuals across conditions. In its standard formulation, however, scGen requires observations from the relevant combinations of conditions in order to estimate the latent shift, which in our setting corresponds to combinations that are not observed. As with Oracle-scVI, scGen can instead be supplied with an externally specified mean shift (denoted *δ* in the original work), allowing it to be adapted to transfer across unseen condition combinations.

#### scINSIGHT

scINSIGHT is a matrix factorization method based on strictly linear factors, with an objective that encourages large differences across conditions. To accommodate combinations of conditions, we represent labels in scINSIGHT using a one-hot encoding 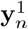. We use *z* and *w* to denote the common and conditional representations, respectively (*W*_*ℓ*1_ and *W*_*ℓ*2_ in the original work).

#### MultiGroupVI

Similar to *Patches*, MultiGroupVI builds on the scVI objective [16] and extends it to model structured conditional variation. In particular, MultiGroupVI introduces group-specific encoders that produce conditional embeddings, together with a mechanism that discourages condition-specific representations from carrying signal when a sample does not originate from the corresponding condition. This heuristic mirrors the formulation adopted by both *Patches* and CSVAE [38], in which the distribution of conditional representations is constrained to collapse toward a point mass at **0** for inactive conditions. In our experiments, we consider MultiGroupVI with one-hot–encoded labels 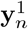 to represent combinations of conditions. For consistency of notation, we denote the common representations by *z* (as in the original work) and the condition-specific representations by *w* (corresponding to *t* in the original formulation).

#### Transfer task setup

To evaluate transfer across conditions, we train scVI, scANVI, scGEN, and *Patches* on a subset of observed condition combinations. Specifically, models are trained using 0DPW (respectively POD7) data from both aged and young (respectively PBS- and verteporfin-treated) mice, together with 4DPW (respectively POD14) data from young (respectively PBS-treated) mice. We then ask whether these models can predict unobserved condition combinations by applying them to cells from 0DPW (respectively POD7) aged (respectively PBS-treated) mice and inferring how their expression profiles would change at 4DPW (respectively POD14) under aged (respectively verteporfin-treated) conditions. An overview of this transfer setup is provided in Supplementary Figure S5.

### Evaluation metrics

We categorized the metrics into two main groups focused on: (1) mixing and separability, and (2) generative accuracy. Mixing and separability metrics quantify the extent to which conditional effects are captured by the models. These metrics were calculated using latent representations – *z* for scVI [16], *z* for scANVI [25] and *z, w* for *Patches*. These metrics were used to evaluate representations for each condition group separately.

Generative accuracy metrics assess the models’ ability to accurately reconstruct or transfer gene expressions of observations across conditions, reflecting model performance. For these metrics, evaluations were conducted on the generated data within the original gene space *g*.

#### Mixing and separability of conditional effects

##### 1. Normalized Condition Average Silhouette Width (cASW)

The silhouette score is a measure of how similar a data point is to its own cluster (within-cluster similarity) compared to other clusters (inter-cluster dissimilarity). To calculate the average silhouette scores across attributes for a given condition group *ζ*^*i*^, we adapt the implementation detailed in [52] - specifically the `silhouette` function - which serves as a wrapper to the original implementation in [44]. The outputted score is a continuous value in the range [0, 1] where values close to 0 indicate non-cluster structure, values around 0.5 indicate overlapping labels and scores closer to 1 indicates compact, distinct clusters.

##### 2. KNN Classification Accuracy (KNN)

Using latent representations of cells and euclidean distance as the distance metric, we train a KNN classifier for a given condition group *ζ*_*i*_ on the latent representations to predict the attributes of each observation. Given attribute labels, the classifier assigns the attribute label of the most common attribute among each observation’s closest *k* = 30 neighbors. We calculate the accuracy score of this classifier, which is the number of correct predictions divided by the number of all predictions. The outputted score is a continuous value in the range [0, 1], and is expected to be high if the clusters are more compact, with neighborhoods of each cell belonging to the same label.

##### 3-4. K-Means (3) Normalized Mutual Information and (4) Adjusted Rand Index

Normalized Mutual Information (NMI) and Adjusted Rand Index (ARI) both measure the similarity of two clusterings. For a given condition group *ζ*^*i*^, the original clusterings are naturally present in the data, which are given by the attributes for each observation. We obtain an additional clustering using latent representations by running K-means with default hyperparameters and *K* = *S*_*i*_ determined by the number of attributes for the given *ζ*^*i*^. Using the two clusterings, we calculate the NMI and ARI through implementations provided in [44] - specifically through the use of `normalized_mutual_info_score` and `adjusted_rand_score` functions. The outputted scores are in the range [0, 1] for NMI and [−0.5, 1] for ARI (even though negative values were not observed), where a higher score indicates greater similarity between clusterings.

#### Generative accuracy - Aggregate metrics

Metrics in this section are applied to normalized pseudobulk gene expression profiles [103, 104], which generalize the analysis to cases where a one-to-one correspondence between generated and ground truth cells is unavailable. To create a pseudobulk gene expression profile *x*^𝒫^ ∈ ℝ^*G*^ for a group of cells **x** ∈ ℝ^*N×G*^, we first compute the total gene expression across all cells in the group. The resulting vector is then scaled to a pre-defined library size *ℓ* as follows:

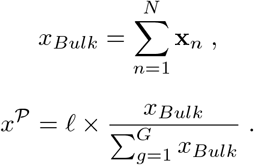

For all applications, we use a default value *ℓ* = 1*e*4. In Figure 5A, we analyze pseudobulk profiles for all cells as well as for specific cell types, comparing these with the pseudobulk profiles derived from reconstructed cells. In Figure 5B we analyze pseudobulk profiles of cell types from the target timepoint. We compare these with transferred profiles of cells mapped from the source timepoint. For simplicity, we use *x* ∈ ℝ^*G*^ to represent the ground-truth pseudobulk gene expression profile and *x*^*′*^ ∈ ℝ^*G*^ for the pseudobulk profile generated by the model.

##### 1. Root Mean Squared Error (RMSE)

The Root mean squared error (RMSE) is a commonly used metric to assess the closeness of two vectors. We report the RMSE between ground truth and generated pseudobulk gene expression profiles. We define RMSE as:

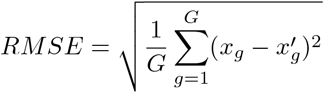

##### 2. Pearson Correlation

Similar to RMSE, Pearson correlation is a widely used metric for comparing gene expression values. Bounded between −1 and 1, it is often preferred over RMSE because of its interpretability. We compute the Pearson correlation coefficient as:

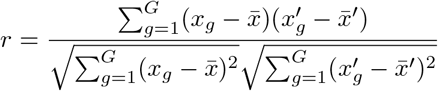

#### Generative accuracy - Point cloud metrics

The metrics in this section are directly applicable to point clouds and, therefore, do not require any matching. This allows them to be used directly with generated observations without additional processing, unlike approaches such as generating pseudobulk expression profiles. In this context, we denote 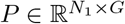 as the point cloud for observations and 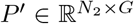 as the point cloud generated by the model, with respective sizes *N*_1_, *N*_2_. For reconstruction tasks, we have *N*_1_ = *N*_2_; however, in transfer tasks, this equality does not necessarily hold. Indeed, for the real datasets considered in this study, it is always the case that *N*_1_≠ *N*_2_.

##### 1. Chamfer Discrepancy (CD)

Chamfer discrepancy (CD) is a metric that quantifies the difference between two point clouds. In our case, these point clouds correspond to observations *P* and generated profiles *P* ^*′*^. CD is defined as:

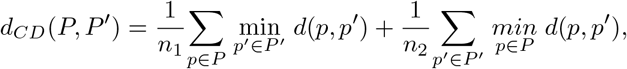

where *d*(*p, q*) is the Euclidean distance between points *p, q*. Each summand iterates over all points in a single point cloud, picks the closest points (supports) from the other cloud, and calculates the sum of distances divided by the number of points. This metric takes values in the range [0, ∞), with lower values indicating higher similarity. However, it is important to note that Chamfer Discrepancy (CD) may lack robustness in certain scenarios. For instance, point clouds with significant differences can sometimes yield low CD values due to the metric being disproportionately influenced by sparsely located points near the boundaries of the point clouds[105, 106].

##### 2. 2-Sliced Wasserstein Distance (SW_2_)

Sliced Wasserstein Distance (*SW*_2_) is another metric used to quantify the difference between two point clouds. We use the implementation provided in [107], where for two probability measures *µ, ν* corresponding to observations *P* and generated points *P* ^*′*^ respectively, the *SW*_2_ is calculated for as:

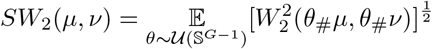

where 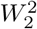 is the 2-Wasserstein distance [105], while *θ*_#_*µ* and *θ*_#_*ν* stand for the pushforwards of the projections *P* ↦⟨*θ, P* ⟩, *P*^*′*^ ↦ ⟨*θ, P*^*′*^⟩ respectively.

### Reproducibility, applicability, robustness, and scalability of *Patches*

#### Reproducibility and applicability

In addition to providing the code used in the study, we offer a high-level workflow API built over `anndata` [102] and a low-level developer API. These both facilitate the reproducibility of our results and streamline the application of conditional generative models to other datasets. We showcase the usage of these APIs through various tutorials, most of which aim to reproduce the behaviour of *Patches* using the simulated and real scRNA-seq datasets featured in this study.

#### Robustness

To ensure the robustness of the generative results, we perform multiple repetitions of training and evaluation. Specifically, we repeat the training process for each benchmark 5 times with different random seeds and repeat the generative processes 10 times across each evaluation before reporting the average performance for a single training process. Tables in Figure 5A,B report the averages of these mean values across 5 training repetitions. For the interpretability module, we investigate 3 distinct model configurations for each data setup: (1) using the standard implementation in [19], (2) incorporating an intercept to the standard implementation and (3) adding an additional ℒ_1_ term to the loss function for conditional sparsity.

#### Scalability

To characterize the computational costs of the deep learning methods considered in this study, we compare runtimes in Supplementary Figure S9A. Relative to other approaches, *Patches* incurs additional runtime, reflecting the extra optimization step introduced by the adversarial classifier, which requires an additional pass through the model. Memory usage is otherwise comparable across methods. For the Vu dataset tutorial, a single pass through the training set (*N* = 21,573 cells) requires 1342 MiB (~1.4 GB) when using default hyperparameters (*Implementation details and hyperparameters*). All experiments were run on a single H100 GPU.

We also observe that, particularly for smaller datasets, *Patches* typically requires more training epochs to reach convergence than other scVI variants, consistent with its more structured training procedure. In addition, we compare two strategies for encoding conditional labels: one-hot encoding over combinations of attributes and a factorized encoding obtained by concatenating individual attributes. Supplementary Figure S9B summarizes the resulting label dimensionality: for *C* = 3 condition groups with *S*_*c*_ attributes per group, the dimensionality of **y**_*n*_ under one-hot encoding of combinations scales exponentially, 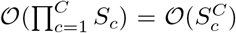 for fixed *S*_*c*_, whereas the factorized representation scales additively, 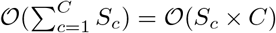. The number of dimensions quickly diverge beyond *S*_*c*_ = 2 for one-hot on combinations, indicating the need for our factorization of **y**_*n*_.

### Analysis of common and cross-condition gene expression modules

Here, we outline the details for comparing the attribute-specific scores derived from the interpretability module of *Patches* with other commonly used quantitative and qualitative approaches for analyzing scRNA-seq datasets.

#### Differential gene expression analysis

For Figure 6A, we obtain significance values for genes using the `rank_genes_groups` function implemented in [13] with default parameters. We perform differential expression testing for each gene by grouping the observations across attributes through the `groupby` parameter, and report the −*log*10 adjusted P-values as significance scores. For rare cases where the adjusted P-value is 0 due to precision constraints, we set the value to 10^−300^.

#### Gene ontology enrichment analysis

GO enrichment analysis was conducted using the `enrichr` wrapper provided in [108, 63] using the gene set `GO_Biological_Process 2023` for mouse. Attribute-specific scores obtained from the interpretability module were ranked and the union of top 10% genes across all attributes were used to construct the input gene list for enrichment testing. Significantly enriched (adjusted P-value *<* 0.05) GO terms were ranked and manually curated to remove redundancy.

## Data availability

Raw data files for the Vu dataset [4] can be accessed through the Gene Expression Omnibus (GEO), under the accession number GSE188432. Raw files for the Mascharak [48] dataset can be found under the accession number GSE186527. The processed data for the in-house snRNA-Seq data [67] are available on GitHub. Additionally, we provide the exact `anndata` objects used in this study through the implemented package.

## Code availability

Implementation of *Patches*, as well as the code base that can be used to reproduce results and apply the models considered in this study to other datasets can be found at: https://github.com/Computational-Morphogenomics-Group/Ladder/tree/patches. Documentation and tutorials on how to apply the models to new datasets will be made publicly available under https://ladder.readthedocs.io/.

## Author contributions

Funding acquisition: BD, YW

Conceptualization: BD, YW

Methodology: OB, BD

Supplementary Data Acquisition: SV, JC

Investigation: OB, SV, MT, DA, JPN, MR, YW, BD

Visualization: OB, BD

Project administration: BD

Supervision: BD

Writing – original draft: OB, BD

Writing – review and editing: OB, DA, MT, MDR YW, BD

## Acknowledgments

BD acknowledges the support of the CIFAR MacMillan Multiscale Human Project. BD and YW acknowledge the support of the Data Science Institute Seed Grant, from Columbia University and NIH grant R00GM140262.

## Supplementary Tables and Figures

**Supplementary Table 1:** Summary of gene level statistics used by Wang et al. [46].

| Score | Type / Condition | Description |
| --- | --- | --- |
| tF | Type | F-statistic from differential gene expression analysis between conditions |
| sPBDS | Condition | Minimum p-value across clusters from pseudobulk-based DSA |
| tPVE / sPVE | Type / Condition | Fraction of variance attributable to conditions |
| HVG | standard gene selection | Gene variance |
| random | control | Drawn uniformly at random |

**Supplementary Table 2:**
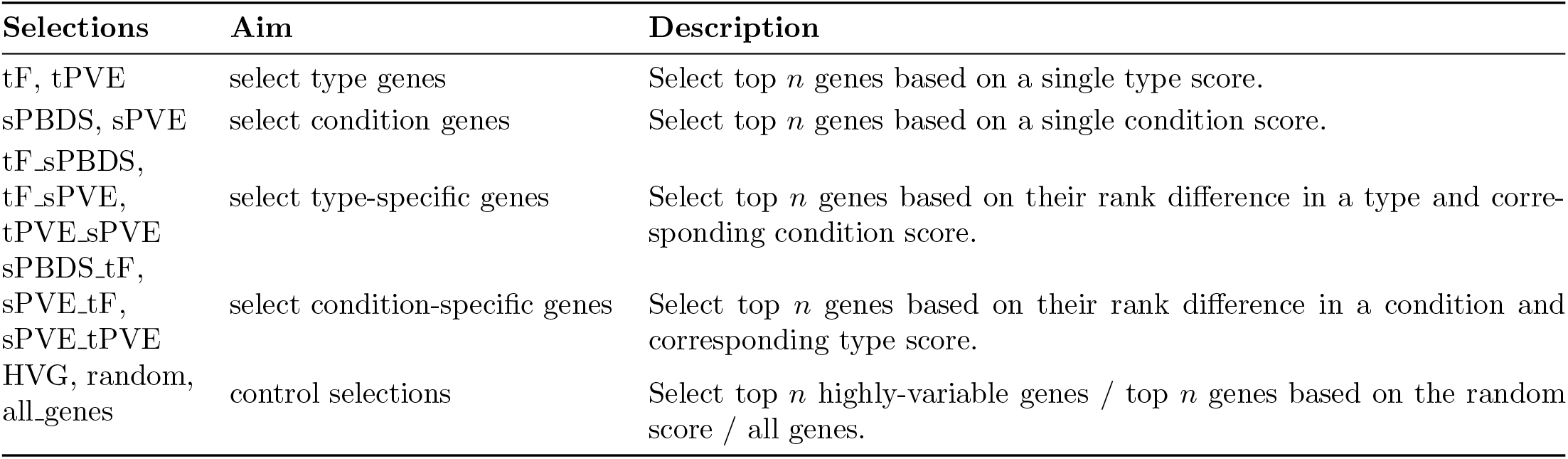
Overview of selection strategies based on type and condition scores for Wang et al. [46].

**Supplementary Table 3:** Overview of cell type and condition scores derived from *Patches*.

| Score | Type / Condition | Description |
| --- | --- | --- |
| cell type scores | Type | *Patches* attribute scores for cell types. |
| condition scores | Condition | *Patches* attribute scores for conditions. |
| tMAV / sMAV | Type / Condition | Maximum absolute value of *Patches* cell type / condition scores. |
| tEN / sEN | Type / Condition | Euclidean norm of *Patches* cell type / condition scores. |
| tMAD / sAD | Type / Condition | Mean absolute difference between *Patches* cell type / condition scores. |

**Supplementary Table 4:** Overview of selection strategies based on scores derived from *Patches*.

| Selections | Aim | Description |
| --- | --- | --- |
| tMAV, tEN, tMAD, | select type genes | Select top $n$ genes based on a single type score. |
| sMAV, sEN, sAD | select condition genes | Select top $n$ genes based on a single condition score. |
| tMAV_sMAV,<br>tEN_sEN,<br>tMAD_sAD | select type-specific genes | Select top $n$ genes based on their rank difference in a type and corresponding condition score. |
| sMAV_tMAV,<br>sEN_tEN,<br>sAD_tMAD | select condition-specific genes | Select top $n$ genes based on their rank difference in a condition and corresponding type score. |

**Supplementary Figure S1:**
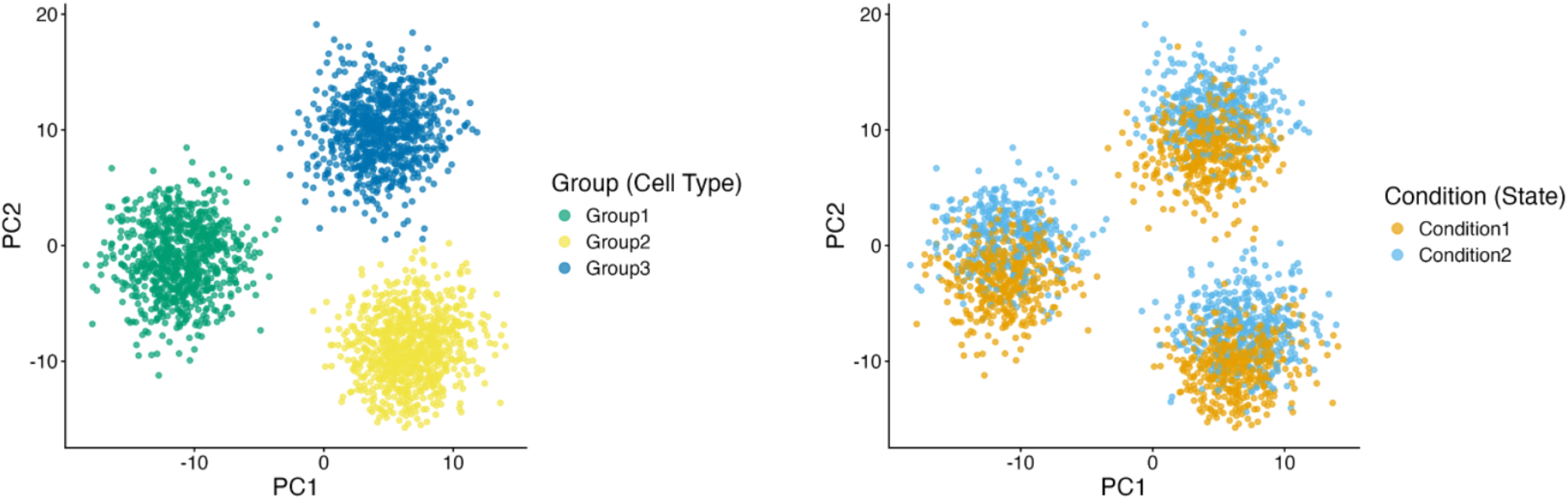
UMAP visualization of the simulated dataset. Cells are colored by cell type (left) and condition (right).

**Supplementary Figure S2:**
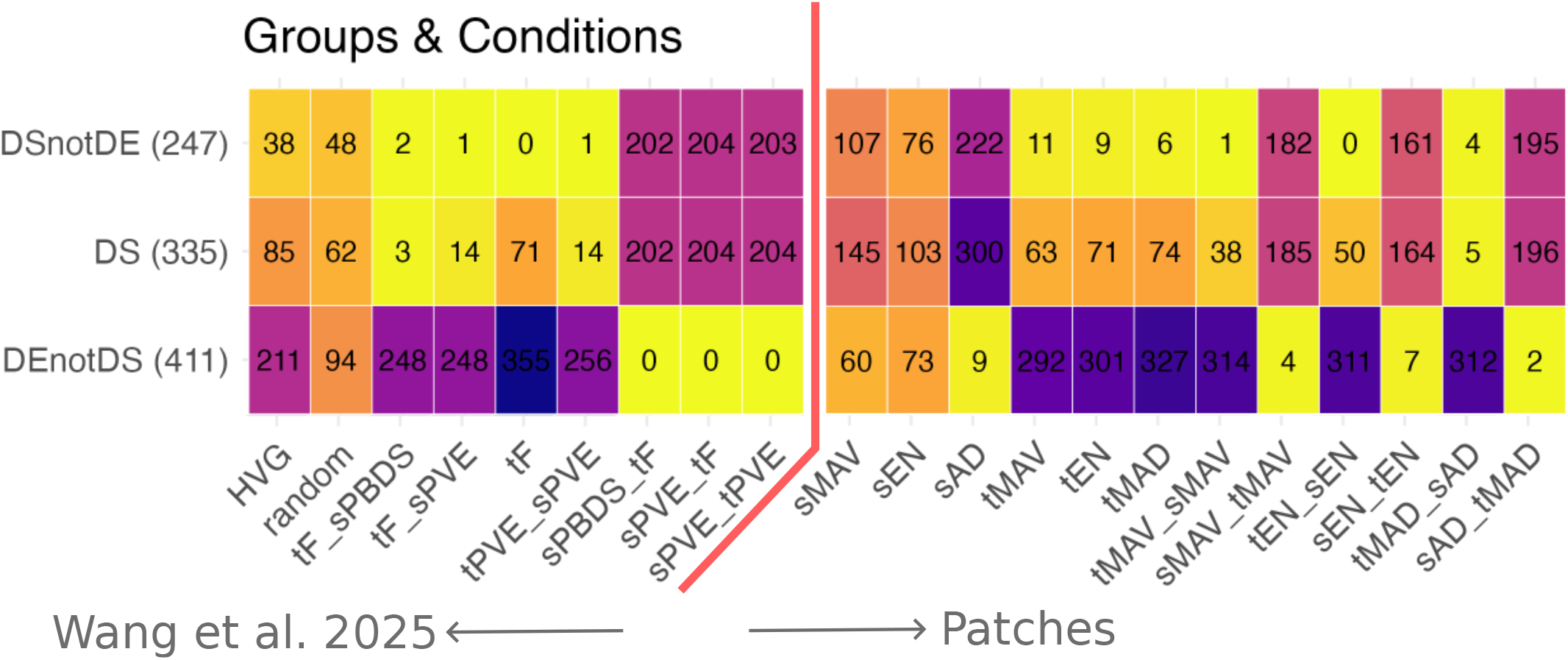
Results from the simulated experiment. Metrics obtained on the embeddings of *Patches* recapitulate the performance of Wang et al. [46] when both cell type and condition labels are provided. DSnotDE denotes recovery for genes that are differentially expressed only across conditions. DS denotes recovery for ground truth conditional genes. DEnotDS denotes recovery for ground truth type genes. This results in identification of gene groups that are discriminative exclusively for either conditions or types, as obtained by both *Patches* and Wang et al. [46].

**Supplementary Figure S3:**
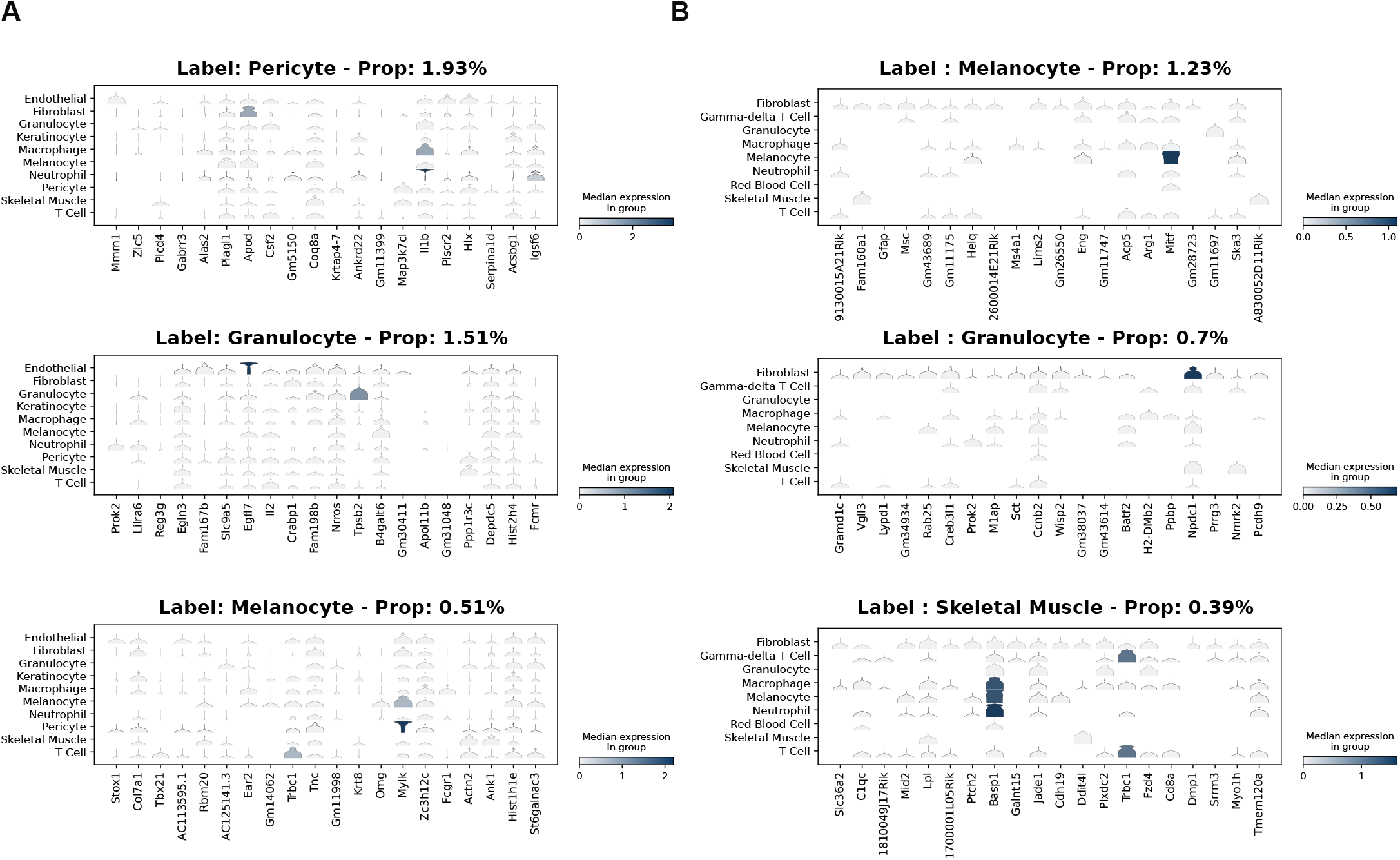
Interpretable *Patches* with less represented cell types. **A, B**, Top 20 marker genes associated with the provided cell type label annotations for the 3 minority cell types in the Vu (**A**) and Mascharak (**B**) datasets, obtained from the linear decoder of Patches. Violin plots show gene expression distributions across all annotated cell types. Compared to majority cell types, *Patches*’ interpretability module appears to be significantly less informative for minority cell types, reemphasizing the fact that deep learning methods require larger number of samples to perform well.

**Supplementary Figure S4:**
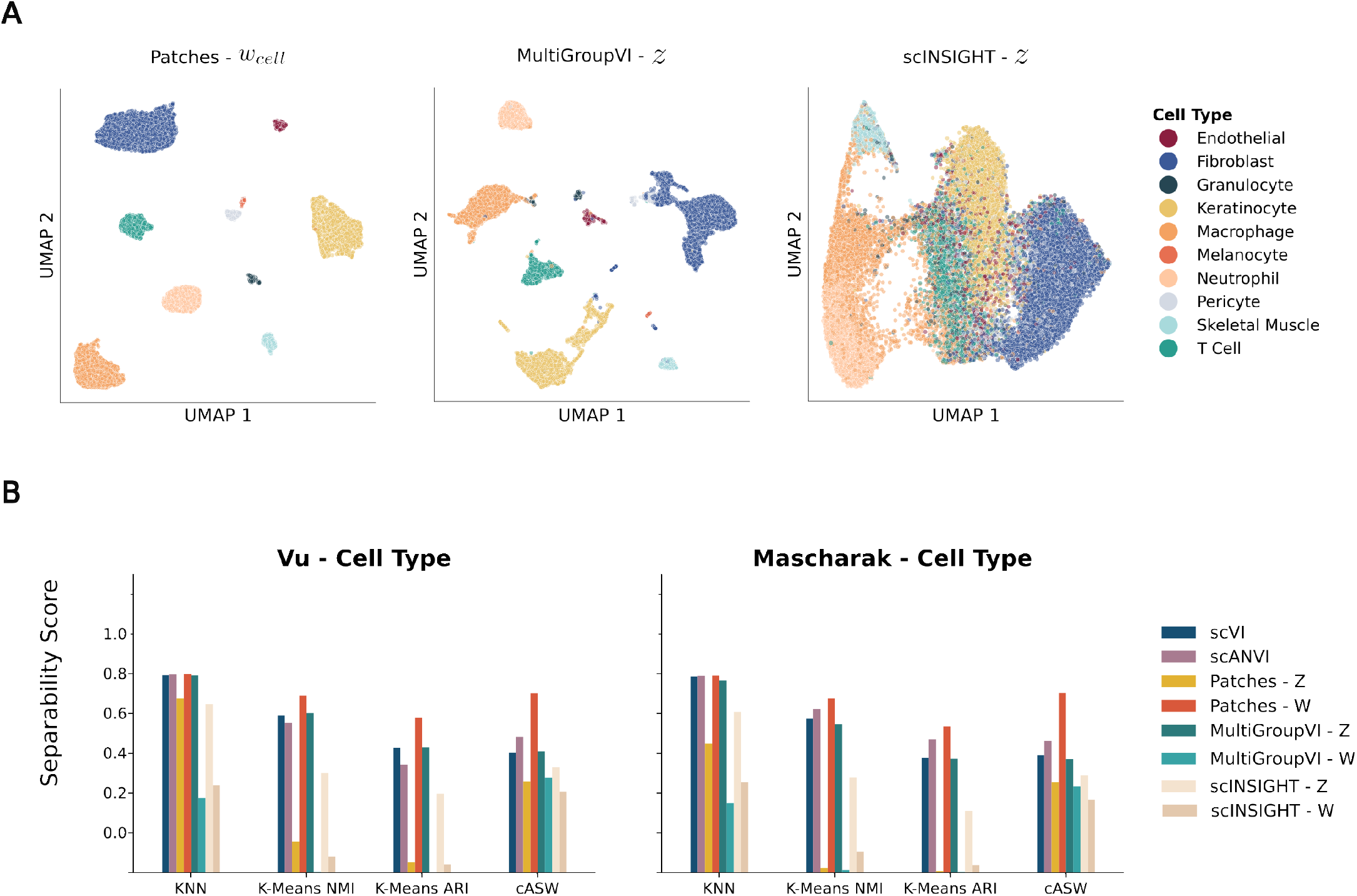
Preservation of Cell Type Information. **A** UMAPs showing cell type specific information is encoded in the conditional embedding of *Patches* (left), and common embeddings of MultiGroupVI (middle) and scINSIGHT (right) for the Vu dataset. **B** Separability scores calculated for cell type on latent representations obtained by running scVI, scANVI, MultiGroupVI, scINSIGHT and Patches on the Vu (left) and Mascharak (right) datasets.

**Supplementary Figure S5:**
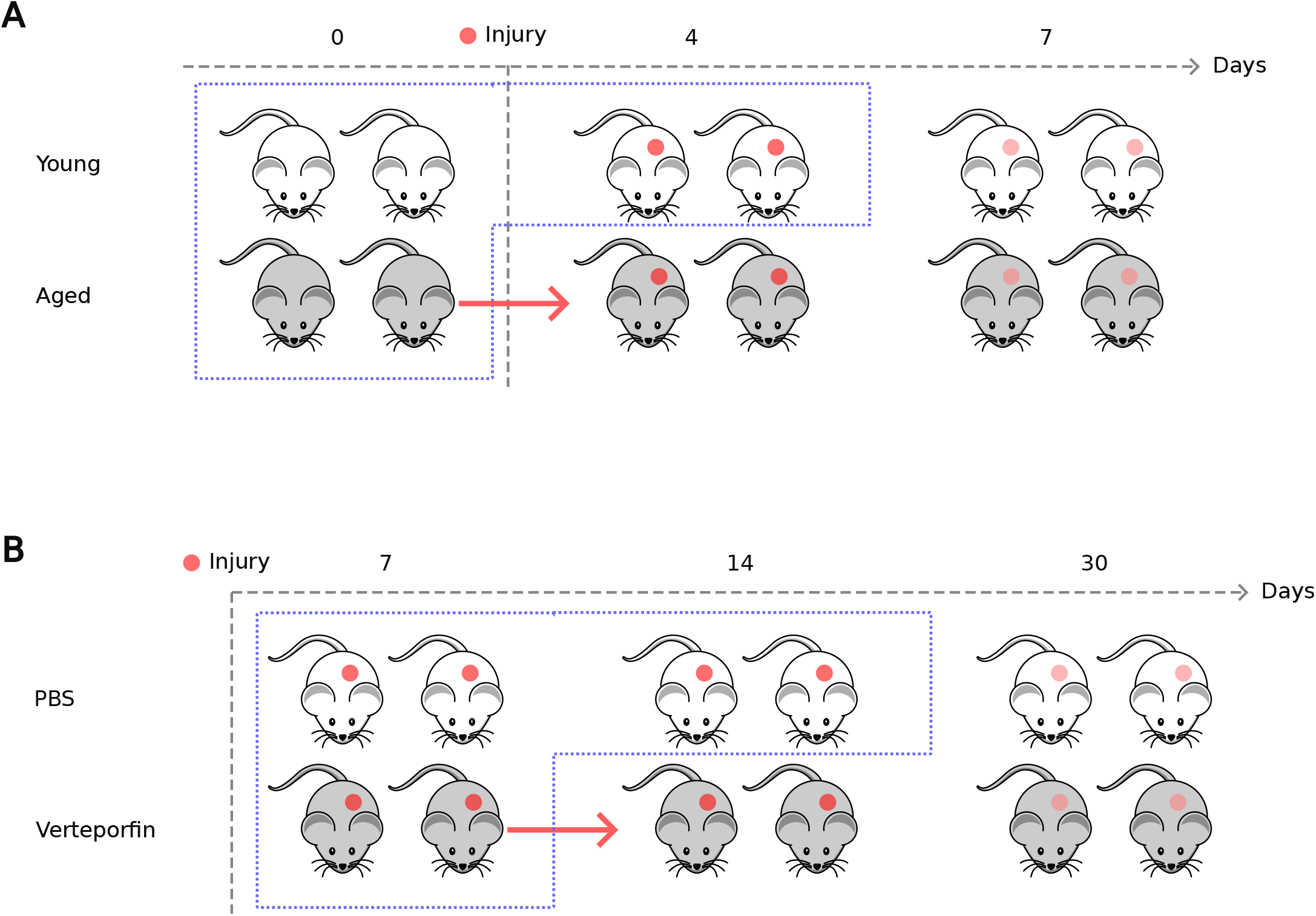
Transfer task setup. **A, B** Models train on the first time point for both aging (**A**) / treatment **B** conditions, and the second time point for only one of aging / treatment conditions (denoted by the dashed blue box). Model performance is evaluated for predicting counterfactuals using cells from the first time point to generate the unobserved aging / treatment condition in the next time point (denoted by the red arrow).

**Supplementary Figure S6:**
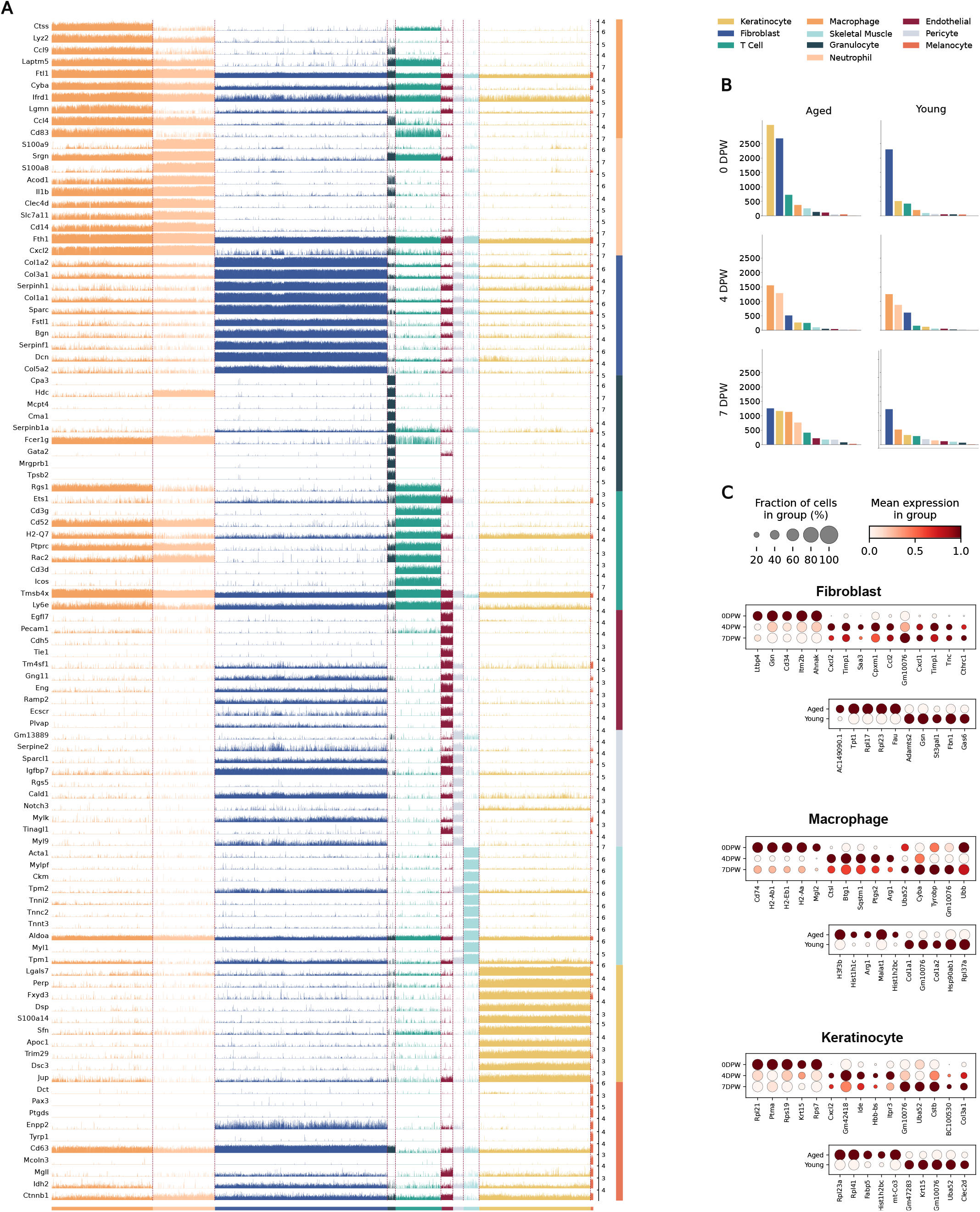
Differential gene expression analysis for the Vu dataset. **A** Track plot showing count distribution across cell types for differentially expressed genes. **B** Sample size distribution across different condition combinations for the Vu dataset. **C** Top differentially expressed genes across time and age for the majority cell types in the Vu dataset.

**Supplementary Figure S7:**
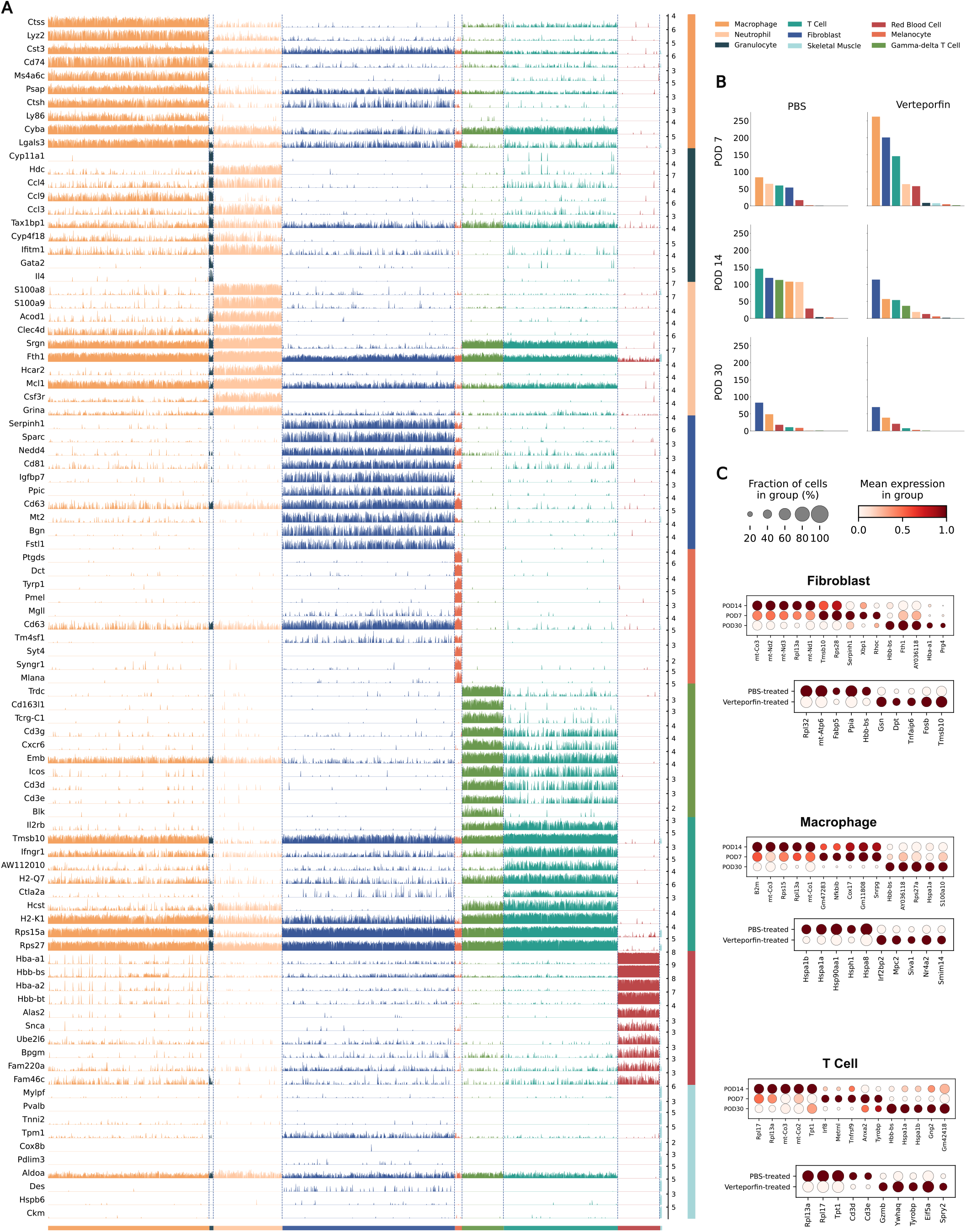
Differential gene expression analysis for the Mascharak dataset. **A** Track plot showing count distribution across cell types for differentially expressed genes. **B** Sample size distribution across different condition combinations for the Mascharak dataset. **C** Top differentially expressed genes across time and treatment for the majority cell types in the Mascharak dataset.

**Supplementary Figure S8:**
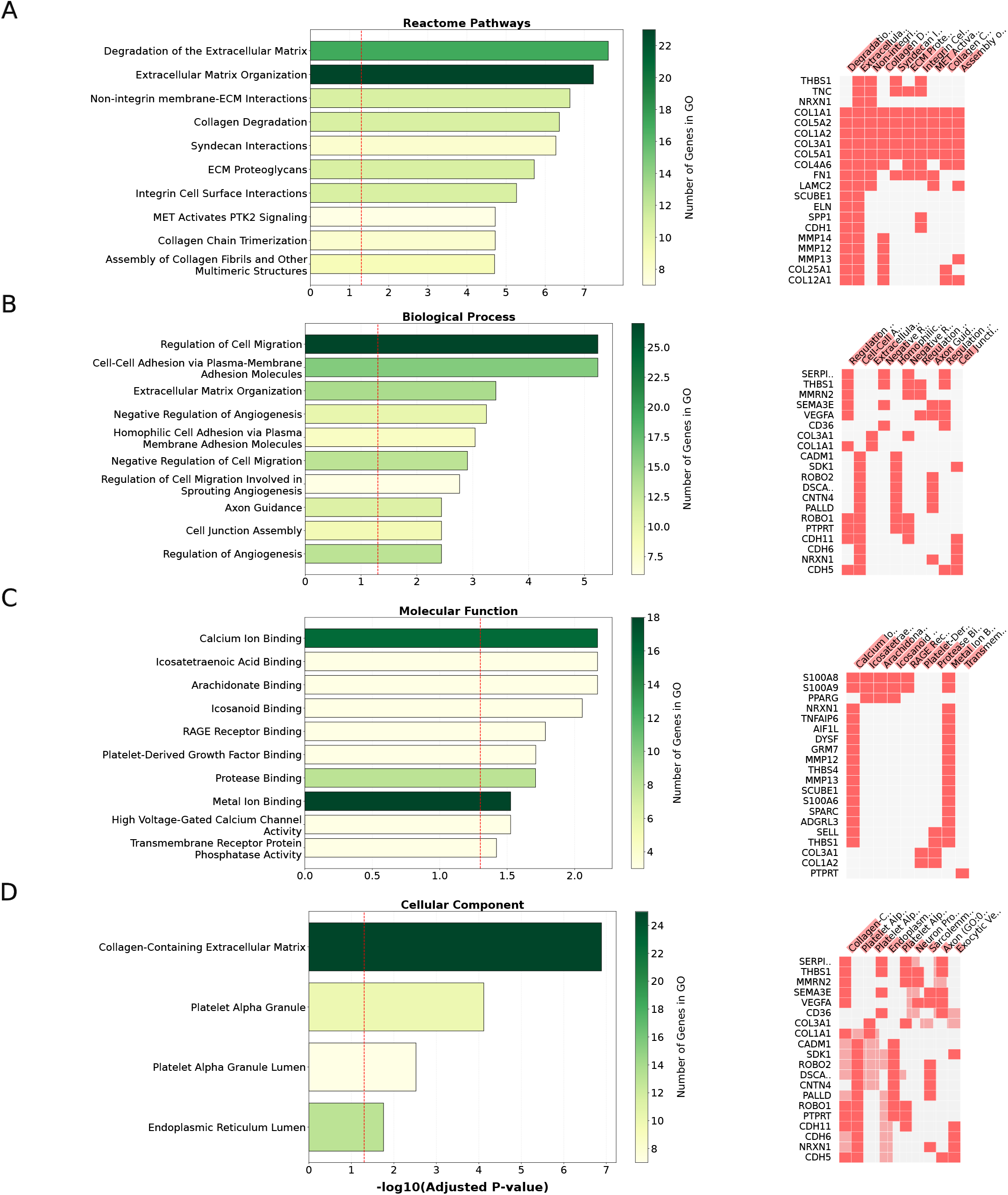
Gene ontology results for the snRNA-seq dataset. GO enrichment analyses are conducted and top GOs with significant (Adjusted *P <* 0.05) enrichment for DPW7 are displayed for the following annotations: **A** Reactome Pathways **B** GO - Biological Process **C** GO - Molecular Function and **D** GO - Cellular Component. Red dashed line denotes significance boundary (−log 0.05). Right column denotes genes associated with GOs on the same row.

**Supplementary Figure S9:**
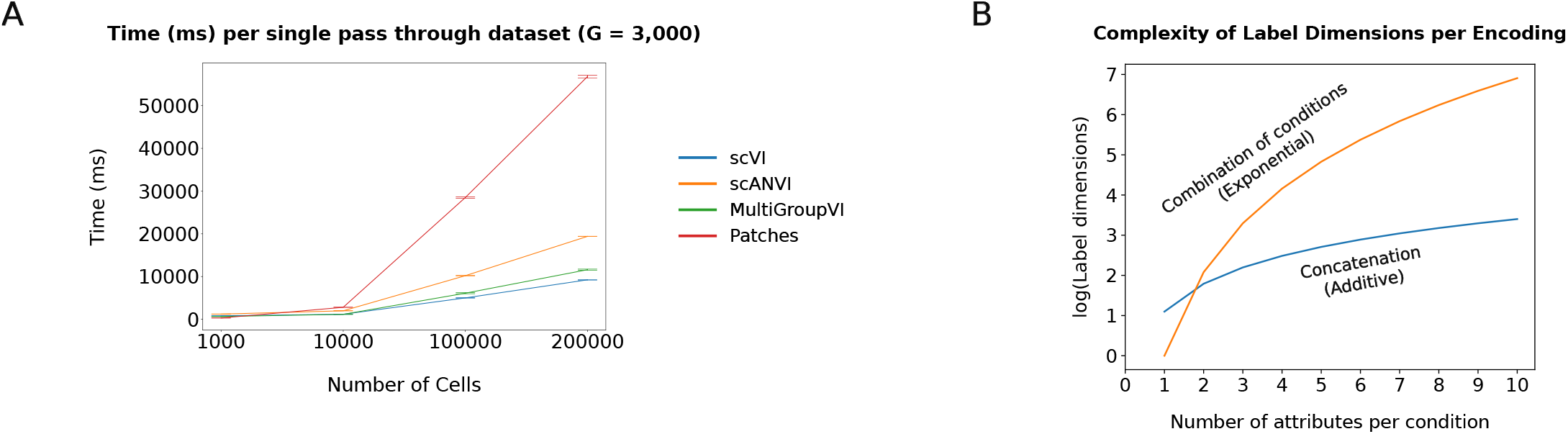
Extended scalability metrics. **A** Time it takes for a single pass through the dataset (in milliseconds) for fixed number of genes *G* = 3, 000 across different number of cells. **B** Complexity of label sizes when considering combinations as one-hot encodings versus concatenation of separate attributes.

## Notes

https://github.com/Computational-Morphogenomics-Group/Ladder/tree/patches

### Competing Interest Statement

The authors have declared no competing interest.

### Summary of Updates

Expanded sections; updated authors; updated supplamentary information

## References

[1] Gonzales, K. A. U. & Fuchs, E. Skin and its regenerative powers: an alliance between stem cells and their niche. Developmental cell 43, 387–401 (2017).

[2] Takeo, M., Lee, W. & Ito, M. Wound healing and skin regeneration. Cold Spring Harbor perspectives in medicine 5, a023267 (2015).

[3] Eming, S. A., Martin, P. & Tomic-Canic, M. Wound repair and regeneration: mechanisms, signaling, and translation. Science translational medicine 6, 265sr6–265sr6 (2014).

[4] Vu, R. et al. Wound healing in aged skin exhibits systems-level alterations in cellular composition and cell-cell communication. Cell Reports 40, 111155 (2022). URL 10.1016/j.celrep.2022.111155.

[5] Haensel, D. et al. Defining epidermal basal cell states during skin homeostasis and wound healing using single-cell transcriptomics. Cell reports 30, 3932–3947 (2020).

[6] Foster, D. S. et al. Integrated spatial multiomics reveals fibroblast fate during tissue repair. Proceedings of the National Academy of Sciences 118, e2110025118 (2021).

[7] Guerrero-Juarez, C. F. et al. Single-cell analysis reveals fibroblast heterogeneity and myeloid-derived adipocyte progenitors in murine skin wounds. Nature communications 10, 650 (2019).

[8] Hu, K. H. et al. Transcriptional space-time mapping identifies concerted immune and stromal cell patterns and gene programs in wound healing and cancer. Cell stem cell 30, 885–903 (2023).

[9] Liu, Z. et al. Spatiotemporal single-cell roadmap of human skin wound healing. bioRxiv (2024). URL 10.1101/2024.08.30.609923.

[10] Almet, A. A., Liu, Y., Nie, Q. & Plikus, M. V. Integrated single-cell analysis reveals spatially and temporally dynamic heterogeneity in fibroblast states during wound healing. Journal of Investigative Dermatology (2024). URL 10.1016/j.jid.2024.06.1281.

[11] Chen, J. C. & Christiano, A. M. Out of many, one: Computational reconstruction of mouse skin using single-cell transcriptomics. Cell Stem Cell 19, 421–422 (2016).

[12] Hao, Y. et al. Dictionary learning for integrative, multimodal and scalable single-cell analysis. Nature Biotechnology 42, 293–304 (2023). URL 10.1038/s41587-023-01767-y.

[13] Wolf, F. A., Angerer, P. & Theis, F. J. SCANPY: large-scale single-cell gene expression data analysis. Genome Biology 19 (2018). URL 10.1186/s13059-017-1382-0.

[14] Kiselev, V. Y., Andrews, T. S. & Hemberg, M. Challenges in unsupervised clustering of single-cell RNA-seq data. Nature Reviews Genetics 20, 273–282 (2019). URL 10.1038/s41576-018-0088-9.

[15] Korsunsky, I. et al. Fast, sensitive and accurate integration of single-cell data with Harmony. Nature methods 16, 1289–1296 (2019).

[16] Lopez, R., Regier, J., Cole, M. B., Jordan, M. I. & Yosef, N. Deep generative modeling for single-cell transcriptomics. Nature Methods 15, 1053–1058 (2018). URL 10.1038/s41592-018-0229-2.

[17] Elyanow, R., Dumitrascu, B., Engelhardt, B. E. & Raphael, B. J. netNMF-sc: leveraging gene–gene interactions for imputation and dimensionality reduction in single-cell expression analysis. Genome Research 30, 195–204 (2020). URL 10.1101/gr.251603.119.

[18] Kunes, R. Z., Walle, T., Land, M., Nawy, T. & Pe’er, D. Supervised discovery of interpretable gene programs from single-cell data. Nature Biotechnology 42, 1084–1095 (2023). URL 10.1038/s41587-023-01940-3.

[19] Svensson, V., Gayoso, A., Yosef, N. & Pachter, L. Interpretable factor models of single-cell RNA-seq via variational autoencoders. Bioinformatics 36, 3418–3421 (2020). URL 10.1093/bioinformatics/btaa169. https://academic.oup.com/bioinformatics/article-pdf/36/11/3418/50670669/bioinformatics_36_11_3418.pdf.

[20] Jin, S. et al. Inference and analysis of cell-cell communication using cellchat. Nature Communications 12 (2021). URL 10.1038/s41467-021-21246-9.

[21] Wang, S., Karikomi, M., MacLean, A. L. & Nie, Q. Cell lineage and communication network inference via opti-mization for single-cell transcriptomics. Nucleic Acids Research 47, e66–e66 (2019). URL 10.1093/nar/gkz204.

[22] Cabello-Aguilar, S. et al. SingleCellSignalR: inference of intercellular networks from single-cell transcriptomics. Nucleic Acids Research 48, e55–e55 (2020). URL 10.1093/nar/gkaa183.

[23] Nazaret, A. et al. Deep generative model deciphers derailed trajectories in acute myeloid leukemia (2023). URL 10.1101/2023.11.11.566719.

[24] Tran, D. et al. Fast and precise single-cell data analysis using a hierarchical autoencoder. Nature Communications 12 (2021). URL 10.1038/s41467-021-21312-2.

[25] Xu, C. et al. Probabilistic harmonization and annotation of single-cell transcriptomics data with deep generative models. Molecular Systems Biology 17 (2021). URL 10.15252/msb.20209620.

[26] Grønbech, C. H. et al. scVAE: variational auto-encoders for single-cell gene expression data. Bioinformatics 36, 4415–4422 (2020). URL 10.1093/bioinformatics/btaa293.

[27] Ashuach, T. et al. Multivi: deep generative model for the integration of multimodal data. Nature Methods 20, 1222–1231 (2023). URL 10.1038/s41592-023-01909-9.

[28] Gayoso, A. et al. Joint probabilistic modeling of single-cell multi-omic data with totalVI. Nature Methods 18, 272–282 (2021). URL 10.1038/s41592-020-01050-x.

[29] Lotfollahi, M., Wolf, F. A. & Theis, F. J. scGen predicts single-cell perturbation responses. Nature Methods 16, 715–721 (2019). URL 10.1038/s41592-019-0494-8.

[30] Lotfollahi, M., Naghipourfar, M., Theis, F. J. & Wolf, F. A. Conditional out-of-distribution generation for unpaired data using transfer VAE. Bioinformatics 36, i610–i617 (2020). URL 10.1093/bioinformatics/btaa800.

[31] Boyeau, P. et al. Deep generative modeling of sample-level heterogeneity in single-cell genomics. bioRxiv (2024). URL https://www.biorxiv.org/content/early/2024/05/10/2022.10.04.510898. https://www.biorxiv.org/content/early/2024/05/10/2022.10.04.510898.full.pdf.

[32] Argelaguet, R. et al. Multi-omics factor analysis—a framework for unsupervised integration of multi-omics data sets. Molecular Systems Biology 14 (2018). URL 10.15252/msb.20178124.

[33] Gundersen, G., Dumitrascu, B., Ash, J. T. & Engelhardt, B. E. End-to-end training of deep probabilistic CCA on paired biomedical observations. In Adams, R. P. & Gogate, V. (eds.) Proceedings of The 35th Uncertainty in Artificial Intelligence Conference, vol. 115 of Proceedings of Machine Learning Research, 945–955 (PMLR, 2020). URL https://proceedings.mlr.press/v115/gundersen20a.html.

[34] Weinberger, E., Lin, C. & Lee, S.-I. Isolating salient variations of interest in single-cell data with contrastivevi. Nature Methods 20, 1336–1345 (2023). URL 10.1038/s41592-023-01955-3.

[35] Lin, K. Z. & Zhang, N. R. Quantifying common and distinct information in single-cell multimodal data with Tilted Canonical Correlation Analysis. Proceedings of the National Academy of Sciences 120 (2023). URL 10.1073/pnas.2303647120.

[36] Jin, S., Zhang, L. & Nie, Q. scAI: an unsupervised approach for the integrative analysis of parallel single-cell transcriptomic and epigenomic profiles. Genome Biology 21 (2020). URL 10.1186/s13059-020-1932-8.

[37] Kana, O. et al. Generative modeling of single-cell gene expression for dose-dependent chemical perturbations. Patterns 4, 100817 (2023). URL https://www.sciencedirect.com/science/article/pii/S2666389923001861.

[38] Klys, J., Snell, J. & Zemel, R. Learning latent subspaces in variational autoencoders (2018). URL https://arxiv.org/abs/1812.06190. 1812.06190.

[39] Creswell, A., Mohamied, Y., Sengupta, B. & Bharath, A. A. Adversarial information factorization (2018). URL https://arxiv.org/abs/1711.05175. 1711.05175.

[40] Andrew, G., Arora, R., Bilmes, J. & Livescu, K. Deep canonical correlation analysis. In Dasgupta, S. & McAllester, D. (eds.) Proceedings of the 30th International Conference on Machine Learning, vol. 28 of Proceedings of Machine Learning Research, 1247–1255 (PMLR, Atlanta, Georgia, USA, 2013). URL https://proceedings.mlr.press/v28/andrew13.html.

[41] Risso, D., Perraudeau, F., Gribkova, S., Dudoit, S. & Vert, J.-P. A general and flexible method for signal extraction from single-cell RNA-seq data. Nature Communications 9 (2018). URL 10.1038/s41467-017-02554-5.

[42] Pierson, E. & Yau, C. ZIFA: Dimensionality reduction for zero-inflated single-cell gene expression analysis. Genome biology 16, 1–10 (2015).

[43] Eraslan, G., Simon, L. M., Mircea, M., Mueller, N. S. & Theis, F. J. Single-cell RNA-seq denoising using a deep count autoencoder. Nature Communications 10 (2019). URL 10.1038/s41467-018-07931-2.

[44] Pedregosa, F. et al. Scikit-learn: Machine learning in Python. Journal of Machine Learning Research 12, 2825–2830 (2011).

[45] Marsland, S. Machine Learning: An Algorithmic Perspective, Second Edition (Chapman & Hall/CRC, 2014), 2nd edn.

[46] Wang, J., Crowell, H. L. & Robinson, M. D. On feature selection to disentangle cell type and state transcriptional programs. BMC Genomics 26 (2025). URL 10.1186/s12864-025-12085-9.

[47] Zappia, L., Phipson, B. & Oshlack, A. Splatter: simulation of single-cell rna sequencing data. Genome Biology 18 (2017). URL 10.1186/s13059-017-1305-0.

[48] Mascharak, S. et al. Multi-omic analysis reveals divergent molecular events in scarring and regenerative wound healing. Cell Stem Cell 29, 315–327.e6 (2022). URL 10.1016/j.stem.2021.12.011.

[49] Yang, X., Bam, M., Becker, W., Nagarkatti, P. S. & Nagarkatti, M. Long noncoding RNA AW112010 promotes the differentiation of inflammatory T cells by suppressing IL-10 expression through histone demethylation. The Journal of Immunology 205, 987–993 (2020). URL 10.4049/jimmunol.2000330.

[50] Weinberger, E., Lopez, R., Huetter, J.-C. & Regev, A. Disentangling shared and group-specific variations in single-cell transcriptomics data with multigroupvi. In Knowles, D. A., Mostafavi, S. & Lee, S.-I. (eds.) Proceedings of the 17th Machine Learning in Computational Biology meeting, vol. 200 of Proceedings of Machine Learning Research, 16–32 (PMLR, 2022). URL https://proceedings.mlr.press/v200/weinberger22a.html.

[51] Qian, K., Fu, S., Li, H. & Li, W. V. scinsight for interpreting single-cell gene expression from biologically heterogeneous data. Genome Biology 23 (2022). URL 10.1186/s13059-022-02649-3.

[52] Luecken, M. D. et al. Benchmarking atlas-level data integration in single-cell genomics. Nature Methods 19, 41–50 (2022).

[53] Witke, W. et al. Hemostatic, inflammatory, and fibroblast responses are blunted in mice lacking gelsolin. Cell 81, 41–51 (1995).

[54] Elabd, C. et al. Oxytocin is an age-specific circulating hormone that is necessary for muscle maintenance and regeneration. Nature communications 5, 4082 (2014).

[55] Garel, S., Marín, F., Grosschedl, R. & Charnay, P. Ebf1 controls early cell differentiation in the embryonic striatum. Development 126, 5285–5294 (1999).

[56] Wu, G. et al. Molecular insights of Gas6/TAM in cancer development and therapy. Cell death & disease 8, e2700–e2700 (2017).

[57] Liu, Y., Lyle, S., Yang, Z. & Cotsarelis, G. Keratin 15 promoter targets putative epithelial stem cells in the hair follicle bulge. Journal of Investigative Dermatology 121, 963–968 (2003). URL 10.1046/j.1523-1747.2003.12600.x.

[58] Phan, Q. M. et al. Lef1 expression in fibroblasts maintains developmental potential in adult skin to regenerate wounds. Elife 9, e60066 (2020).

[59] Ge, W. et al. Single-cell transcriptome profiling reveals dermal and epithelial cell fate decisions during embryonic hair follicle development. Theranostics 10, 7581 (2020).

[60] Hughes, T. K. et al. Second-strand synthesis-based massively parallel scRNA-seq reveals cellular states and molecular features of human inflammatory skin pathologies. Immunity 53, 878–894 (2020).

[61] Fu, Z. D., Csanaky, I. L. & Klaassen, C. D. Effects of aging on mRNA profiles for drug-metabolizing enzymes and transporters in livers of male and female mice. Drug Metabolism and Disposition 40, 1216–1225 (2012).

[62] Zhang, K. et al. Ramanomics decodes spatial vibrational-molecular architecture and rewiring in aging and repair. bioRxiv (2025). URL 10.64898/2025.12.04.692337.

[63] Fang, Z., Liu, X. & Peltz, G. GSEApy: a comprehensive package for performing gene set enrichment analysis in Python. Bioinformatics 39 (2022). URL 10.1093/bioinformatics/btac757.

[64] Angelini, A., Trial, J., Ortiz-Urbina, J. & Cieslik, K. A. Mechanosensing dysregulation in the fibroblast: A hallmark of the aging heart. Ageing Research Reviews 63, 101150 (2020). URL 10.1016/j.arr.2020.101150.

[65] Coulombe, P. A., Ma, L., Yamada, S. & Wawersik, M. Intermediate filaments at a glance. Journal of cell science 114, 4345–4347 (2001).

[66] Nishiguchi, M. A., Spencer, C. A., Leung, D. H. & Leung, T. H. Aging suppresses skin-derived circulating sdf1 to promote full-thickness tissue regeneration. Cell Reports 24, 3383–3392.e5 (2018). URL 10.1016/j.celrep.2018.08.054.

[67] Cheong, J. C., Van Deursen, S., Amador, D., Hiner, S. & Woappi, Y. 4d multimodal wound healing atlas reveals organ-level controls of repair phase transitions. bioRxiv (2026). URL https://www.biorxiv.org/content/early/2026/01/16/2026.01.15.699736. https://www.biorxiv.org/content/early/2026/01/16/2026.01.15.699736.full.pdf.

[68] Wu, Y. et al. Scube3 is an endogenous TGF-β receptor ligand and regulates the epithelial-mesenchymal transition in lung cancer. Oncogene 30, 3682–3693 (2011).

[69] Lorenzo, P., Bayliss, M. T. & Heinegård, D. A novel cartilage protein (CILP) present in the mid-zone of human articular cartilage increases with age. Journal of Biological Chemistry 273, 23463–23468 (1998).

[70] van Nieuwenhoven, F. A. et al. Cartilage intermediate layer protein 1 (CILP1): a novel mediator of cardiac extracellular matrix remodelling. Scientific reports 7, 16042 (2017).

[71] Vannahme, C. et al. Characterization of SMOC-1, a novel modular calcium-binding protein in basement membranes. Journal of Biological Chemistry 277, 37977–37986 (2002).

[72] Mahley, R. W. & Huang, Y. Apolipoprotein E: from atherosclerosis to Alzheimer’s disease and beyond. Current opinion in lipidology 10, 207–218 (1999).

[73] Sun, J. et al. Single-cell RNA sequencing reveals dysregulation of spinal cord cell types in a severe spinal muscular atrophy mouse model. PLoS Genetics 18, e1010392 (2022).

[74] Visscher, M. O. et al. Newborn infant skin gene expression: Remarkable differences versus adults. PLoS One 16, e0258554 (2021).

[75] Wu, S. et al. Single-cell transcriptomics reveals lineage trajectory of human scalp hair follicle and informs mechanisms of hair graying. Cell Discovery 8, 49 (2022).

[76] Steinbinder, J., Sachslehner, A. P., Holthaus, K. B. & Eckhart, L. Comparative genomics of monotremes provides insights into the early evolution of mammalian epidermal differentiation genes. Scientific Reports 14, 1437 (2024).

[77] Lee, S.-C. et al. Human trichohyalin gene is clustered with the genes for other epidermal structural proteins and calcium-binding proteins at chromosomal locus 1q21. Journal of Investigative Dermatology 100, 65–68 (1993). URL 10.1111/1523-1747.ep12354504.

[78] Westgate, G. E., Ginger, R. S. & Green, M. R. The biology and genetics of curly hair. Experimental Dermatology 26, 483–490 (2017). URL 10.1111/exd.13347.

[79] Medland, S. E. et al. Common variants in the trichohyalin gene are associated with straight hair in europeans. The American Journal of Human Genetics 85, 750–755 (2009). URL 10.1016/j.ajhg.2009.10.009.

[80] Ogawa, Y., Kawamura, T. & Shimada, S. Zinc and skin biology. Archives of biochemistry and biophysics 611, 113–119 (2016).

[81] Martin, G. et al. The human fatty acid transport protein-1 (SLC27A1; FATP-1) cDNA and gene: organization, chromosomal localization, and expression. Genomics 66, 296–304 (2000).

[82] Mann, G. B. et al. Mice with a null mutation of the tgfα gene have abnormal skin architecture, wavy hair, and curly whiskers and often develop corneal inflammation. Cell 73, 249–261 (1993).

[83] Ponticos, M. et al. Col1a2 enhancer regulates collagen activity during development and in adult tissue repair. Matrix biology 22, 619–628 (2004).

[84] Bennett, R. D., Pittelkow, M. R. & Strehler, E. E. Immunolocalization of the tumor-sensitive calmodulin-like protein CALML3 in normal human skin and hyperproliferative skin disorders. PLoS One 8, e62347 (2013).

[85] Yu, B. & Kumbier, K. Veridical data science. Proceedings of the National Academy of Sciences 117, 3920–3929 (2020). URL 10.1073/pnas.1901326117.

[86] Hiebert, P. R. et al. Granzyme b contributes to extracellular matrix remodeling and skin aging in apolipoprotein e knockout mice. Experimental gerontology 46, 489–499 (2011).

[87] Farage, M. A., Miller, K. W., Elsner, P. & Maibach, H. I. Characteristics of the aging skin. Advances in wound care 2, 5–10 (2013).

[88] Locatello, F. et al. Challenging common assumptions in the unsupervised learning of disentangled representations. In Chaudhuri, K. & Salakhutdinov, R. (eds.) Proceedings of the 36th International Conference on Machine Learning, vol. 97 of Proceedings of Machine Learning Research, 4114–4124 (PMLR, 2019). URL https://proceedings.mlr.press/v97/locatello19a.html.

[89] Abid, A. & Zou, J. Contrastive variational autoencoder enhances salient features (2019). URL https://arxiv.org/abs/1902.04601. 1902.04601.

[90] Steele, L. et al. A single-cell and spatial genomics atlas of human skin fibroblasts reveals shared disease-related fibroblast subtypes across tissues. Nature Immunology 26, 1807–1820 (2025). URL 10.1038/s41590-025-02267-8.

[91] Restrepo, P. et al. A single-cell spatial transcriptomic census of human skin anatomy. bioRxiv (2025). URL 10.1101/2025.09.22.676865.

[92] Lotfollahi, M. et al. Predicting cellular responses to complex perturbations in high-throughput screens. Molecular Systems Biology 19 (2023). URL 10.15252/msb.202211517.

[93] Khemakhem, I., Kingma, D. P., Monti, R. P. & Hyvärinen, A. Variational autoencoders and nonlinear ICA: A unifying framework (2020). URL https://arxiv.org/abs/1907.04809. 1907.04809.

[94] Salzmann, M., Ek, C. H., Urtasun, R. & Darrell, T. Factorized orthogonal latent spaces. In Teh, Y. W. & Tit-terington, M. (eds.) Proceedings of the Thirteenth International Conference on Artificial Intelligence and Statistics, vol. 9 of Proceedings of Machine Learning Research, 701–708 (PMLR, Chia Laguna Resort, Sardinia, Italy, 2010). URL https://proceedings.mlr.press/v9/salzmann10a.html.

[95] Blei, D. M., Kucukelbir, A. & McAuliffe, J. D. Variational inference: A review for statisticians. Journal of the American Statistical Association 112, 859–877 (2017). URL 10.1080/01621459.2017.1285773.

[96] Bingham, E. et al. Pyro: Deep universal probabilistic programming. J. Mach. Learn. Res. 20, 28:1–28:6 (2019). URL http://jmlr.org/papers/v20/18-403.html.

[97] He, K., Zhang, X., Ren, S. & Sun, J. Delving deep into rectifiers: Surpassing human-level performance on ImageNet classification (2015). URL https://arxiv.org/abs/1502.01852. 1502.01852.

[98] Paszke, A. et al. PyTorch: An imperative style, high-performance deep learning library (2019). URL https://arxiv.org/abs/1912.01703. 1912.01703.

[99] Chandra, N. K., Dunson, D. B. & Xu, J. Inferring covariance structure from multiple data sources via subspace factor analysis. Journal of the American Statistical Association 120, 1239–1253 (2024). URL 10.1080/01621459.2024.2408777.

[100] Wolock, S. L., Lopez, R. & Klein, A. M. Scrublet: Computational identification of cell doublets in single-cell transcriptomic data. Cell Systems 8, 281–291.e9 (2019). URL 10.1016/j.cels.2018.11.005.

[101] Bernstein, N. J. et al. Solo: Doublet identification in single-cell RNA-Seq via semi-supervised deep learning. Cell Systems 11, 95–101.e5 (2020). URL 10.1016/j.cels.2020.05.010.

[102] Virshup, I., Rybakov, S., Theis, F. J., Angerer, P. & Wolf, F. A. anndata: Annotated data (2021). URL 10.1101/2021.12.16.473007.

[103] Zhang, R. et al. Cross-species imputation and comparison of single-cell transcriptomic profiles. bioRxiv (2024). URL https://www.biorxiv.org/content/early/2024/08/12/2023.10.19.563173. https://www.biorxiv.org/content/early/2024/08/12/2023.10.19.563173.full.pdf.

[104] Cheng, J. et al. Unraveling the timeline of gene expression: A pseudotemporal trajectory analysis of single-cell rna sequencing data. F1000Research 12, 684 (2023).

[105] Nguyen, T. et al. Point-set distances for learning representations of 3d point clouds. In 2021 IEEE/CVF Interna-tional Conference on Computer Vision (ICCV), 10458–10467 (IEEE Computer Society, Los Alamitos, CA, USA, 2021). URL https://doi.ieeecomputersociety.org/10.1109/ICCV48922.2021.01031.

[106] Achlioptas, P., Diamanti, O., Mitliagkas, I. & Guibas, L. Learning representations and generative models for 3D point clouds (2018). URL https://arxiv.org/abs/1707.02392. 1707.02392.

[107] Flamary, R. et al. POT: Python Optimal Transport. Journal of Machine Learning Research 22, 1–8 (2021). URL http://jmlr.org/papers/v22/20-451.html.

[108] Chen, E. Y. et al. Enrichr: interactive and collaborative HTML5 gene list enrichment analysis tool. BMC Bioin-formatics 14 (2013). URL 10.1186/1471-2105-14-128.

